# Ubiquitination-mediated mitochondrial protein degradation ensures seedling emergence by regulating ER-mitochondrial interaction and mitophagy

**DOI:** 10.64898/2026.03.10.710782

**Authors:** Zhen Tian, Yawen Huo, Chengyang Li, Qiwei Zheng, Fan Hu, Jing Li, Juncai Ma, Xiaolu Qu, Yunjiang Cheng, Byung-Ho Kang, Patrick Duckney, Pengwei Wang

## Abstract

Seedling emergence is a pivotal step of plant survival, requiring rapid hypocotyl elongation for soil penetration. This energy-demanding process necessitates active mitochondrial respiration, which inevitably induces oxidative damage. Therefore, plants evolved a quality control mechanism that selectively removes dysfunctional mitochondria through the mitophagy pathway. Here, we identified SPL2, a mitochondrial E3 ligase which is essential for hypocotyl elongation and seedling emergence through degrading mitochondrial outer membrane proteins, such as TRB1 and FIS1A. Intriguingly, these proteins also interact with an ER protein, VAP27-1, forming a complex at the ER-mitochondria contact sites, which is essential for mitophagy initiation. The *spl2* mutant exhibits enhanced ER-mitochondrial tethering and mitophagy activation, whereas its overexpression has the opposite effects. The expression of SPL2 increases after light perception, in agreement with the reduced mitophagy. Collectively, our findings reveal novel mechanistic insights into seedling emergence, which are coordinated through protein ubiquitination, ER-mitochondrial interaction, and mitophagy.

## Main

The hypocotyl is the primary tissue that is responsible seedling emergence; it elongating rapidly and provides mechanical force to penetrate the soil surface. This is an energetically intensive process dependent on mitochondrial respiration and is therefore sensitive to factors that cause mitochondrial dysfunction, which may affect seed germination, hypocotyl elongation, and seedling viability^1–7^. However, the cellular mechanisms underlying the maintenance of mitochondrial homeostasis during seedling emergence remain largely elusive. Eukaryotes have evolved intricate quality control pathways to sustain healthy mitochondria. One of the best-known examples is mitophagy, which eliminates dysfunctional mitochondria to maintain mitochondrial integrity and cell homeostasis. In addition, mitochondrial activities are influenced by other organelles that stay in close contact, such as the ER network, which links to mitochondria through ER-mitochondrial contact sites (EMCSs)^8,9^. However, the precise regulation of mitochondrial function in response to diverse developmental cues and environmental signals remains unclear in plants.

To further investigate the regulatory mechanisms of plant mitochondrial maintenance, etiolated seedlings during emergence were selected as an experimental model, which undergoes drastic changes in metabolic rate linked to developmental (seedling germination, cell elongation) and environmental cues (responses to light). Furthermore, this experimental system permits clear investigation of mitochondrial metabolism, easily distinguishable from photosynthesis, as etiolated seedlings are non-photosynthetic and growth at this stage primarily relies on stored energy reserves^2^. During hypocotyl elongation, mitochondrial dynamics underwent dramatic changes, accompanied by elevated autophagy levels and increased mitochondrial turnover (Extended Data Fig. 1a-d), in agreement with a previous study using dark grown seedlings^10^. We therefore hypothesise that a regulatory mechanism may exist to coordinate mitophagy with rapid growth. In animals, the PINK1-Parkin module is the most characterised mitophagy pathway, where the E3 ligase Parkin ubiquitinates outer mitochondrial membrane proteins and targets damaged mitochondria for degradation^11,12^. However, no Parkin homologs have been identified in plant genomes, suggesting the potential existence of analogous pathways in plants that may contribute to seedling emergence and hypocotyl elongation. We searched for putative E3 ligases from the published mitochondrial proteome of different plants^13–15^, and found SPL2 as a putative candidate. This protein family includes SP1 and its two homologs, SPL1 and SPL2, are RING-type E3 ligases. SP1 has been shown to localise to chloroplasts and peroxisomes, regulating the degradation of plastid/peroxisome membrane-localised protein import machineries (e.g. TOC complex, PEX13)^16–19^. In contrast, SPL2 is a more distantly related SP1^20^, and its specific functions remain poorly understood. Using the *Nicotiana benthamiana* transient expression system and transgenic Arabidopsis, we found the SPL2-GFP fusion protein predominantly localised to the outer mitochondrial membrane (OMM) in leaf and etiolated hypocotyl cells, and weak plastid labelling was also observed (Fig. 1a, Extended Data Fig. 2a-b). Such localisation was further confirmed using super-resolution microscopy (SIM), where SPL2-GFP was co-expressed with an OMM marker protein, RFP-TOM20-3 (Fig. 1b-c). Furthermore, SPL2-GFP was able to recruit a cytosolic mRFP-NbG (a nanobody against GFP) to the mitochondrial surface, confirming SPL2 is an outer mitochondrial membrane protein with its GFP-tagged domain oriented toward the cytoplasm (Extended Data Fig. 2c-d).

**Fig 1.**
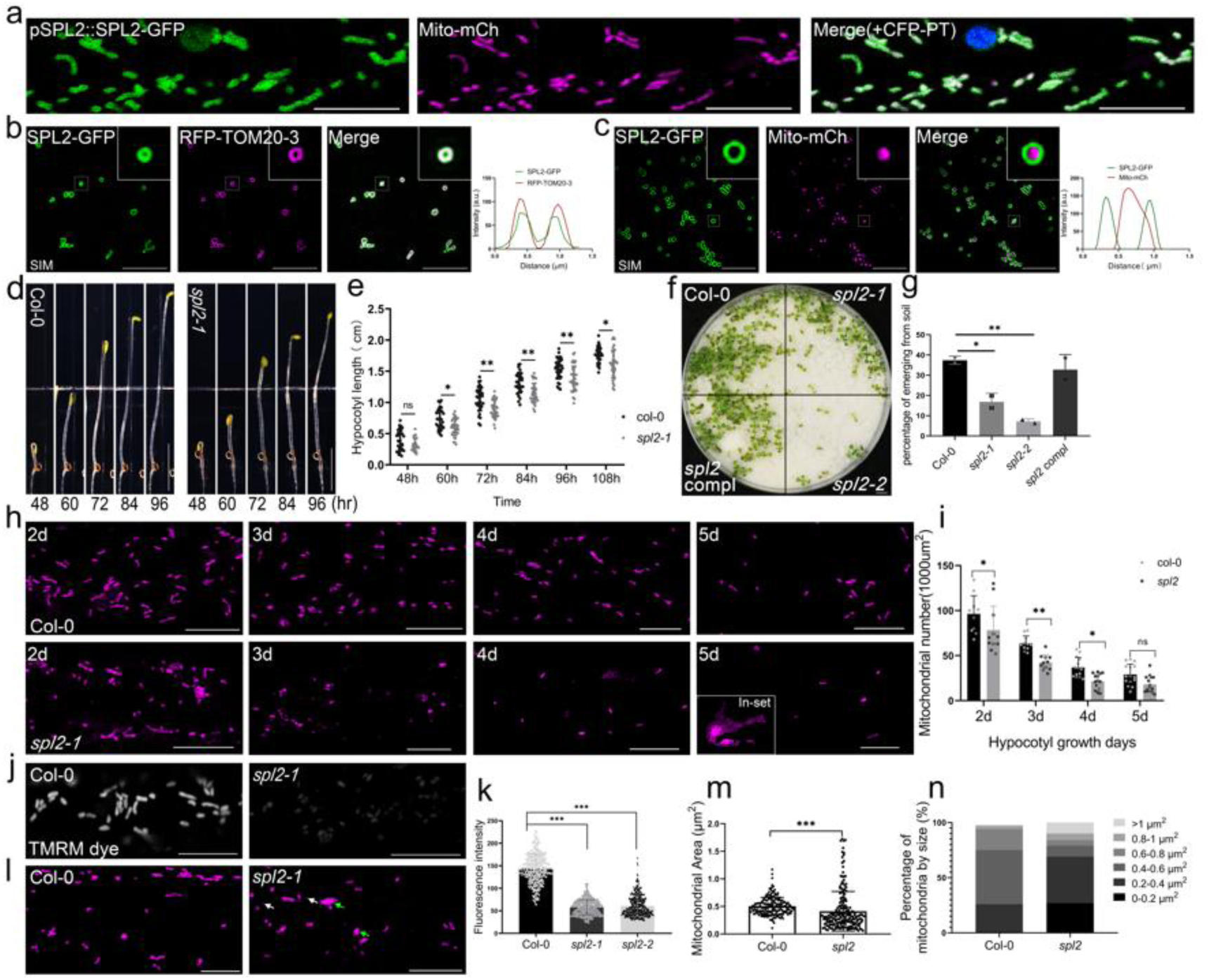
The mitochondrial localised SPL2 mediates seedling emergence. **a.** pSPL2::SPL2-GFP is predominantly targeted to mitochondria in Arabidopsis hypocotyl cells (marked by Mito-mCherry, magenta), and some weak plastid labelling is also identified (using CFP-PT, blue). Scale bar = 10 μm**. b-c.** Super-resolution microscopy (SIM) indicates that SPL2-GFP co-localises with the outer mitochondrial membrane marker RFP-TOM20-3 (b) but does not overlap with the matrix marker Mito-mCherry (c) in *N. benthamiana* leaf epidermal cells. Scale bar = 10 μm**. d-e**. Analysis of hypocotyl elongation in dark-grown (etiolated) seedlings, with measurements taken every 12 h, reveals that *spl2-1* mutants exhibit a significantly lower elongation rate than Col-0. Scale bar = 0.5 cm. n≥30. **f-g.** Soil emergence assay of Col-0, *spl2* mutants (*spl2-1*, *spl2-2*), and the complementation line (pSPL2::SPL2-GFP/*spl2-*1). **f.** Representative images of seedlings germinated under a 3 mm sand cover and imaged after 10 days of light exposure. Scale bar = 0.5 cm. **g.** Quantification of soil emergence rates. Expression of pSPL2::SPL2-GFP rescues the *spl2-1* mutant phenotype. n≥100. **h-i**. In the *spl2* mutant hypocotyl cells stably expressing Mito-mCherry, mitochondrial number was significantly altered compared to Col-0. Mitochondrial aggregates (h, inset) were also frequently observed in 5-day-old etiolated *spl2* mutant seedlings. The number of mitochondria was counted in randomly selected 1000 μm² regions from at least 11 cells of three seedlings. Scale bar = 10 μm**. j.** The mitochondrial membrane potential (MMP), assessed by the fluorescent dye tetramethylrhodamine methyl ester (TMRM), was significantly decreased in root cells of 5-day-old *spl2* mutant seedlings compared to Col-0. **k.** Quantification of TMRM fluorescence intensity in cells imaged as in (j). Over 340 mitochondria from at least 30 cells (n ≥300) were used for the quantification. Scale bar = 10 μm**. l-n.** The presence of mitochondrial fragments (diameter < 0.2 μm; white arrow) and aggregates (diameter > 1 μm; green arrow) in the *spl2* mutant hypocotyl cells. Over 200 mitochondria from 10 cells (n ≥200) were used for the quantification. Scale bar = 10 μm. Statistical analyses were performed using a two-tailed Student’s *t*-test for (m), a one-way ANOVA with Dunnett’s multiple comparisons test for (g) and (k), and a two-way ANOVA with Dunnett’s multiple comparisons test for (e) and (i). Error bars represent SD, and asterisks indicate significant differences in means at *P* < 0.05.

We then generated the *spl2* Arabidopsis knock-out mutant using the CRISPR-Cas9 system, and selected two independent lines for phenotype studies (*spl2-1, spl2-2;* Extended Data Fig. 2e). In the *spl2* mutants, seed germination and hypocotyl elongation rates are significantly reduced (Fig. 1d-e, Extended Data Fig. 2f), ultimately affecting seedling emergence (Fig. 1f-g). These phenotypes are likely attributed to mitochondrial abnormalities, including reduced mitochondrial membrane potential and decreased mitochondrial number (Fig. 1h-k). Meanwhile, mitochondrial fragmentation and aggregations are also abundant (Fig. 1h, l-n; Extended Data Fig. 2g-h). Gene complementation assay using pSPL2::SPL2-GFP confirmed that the seedling emergence and hypocotyl phenotype of the *spl2* mutant is caused by SPL2 loss-of-function (Fig. 1f-g; Extended Data Fig. 2i).

To better understand the function of SPL2, we selected several key mitochondrial regulators (see the method section for details) and performed protein-protein interaction assays using the split-ubiquitin yeast two-hybrid system. We identified proteins regulating ER-mitochondrial interaction (e.g., MIRO, TRB1)^9,21^, mitochondrial fission (e.g., FIS1A, PMD2)^22–24^, and mitophagy (e.g., TRB1)^9^ that interact with SPL2 (Extended Data Fig. 3a). To avoid replication, TRB1 and FIS1A were selected for further studies. In *N. benthamiana* leaf cells, SPL2 colocalises and interacts with FIS1A and TRB1 at mitochondria, as indicated by the transient expression and BiFC assay (Fig. 2a-b). These interactions were confirmed using a co-immunoprecipitation assay, in which SPL2-GFP fusion was co-expressed with HA-tagged TRB1 or FIS1A, and these proteins could be pulled down together with SPL2-GFP using the GFP-Trap system (Fig. 2c-d). It should be noted that the FIS1A protein is also found in peroxisomes and chloroplasts according to a previous study^23,25^. Given its predominant mitochondria localisation, we investigated whether it may be a direct regulator of mitochondrial function.

**Fig 2.**
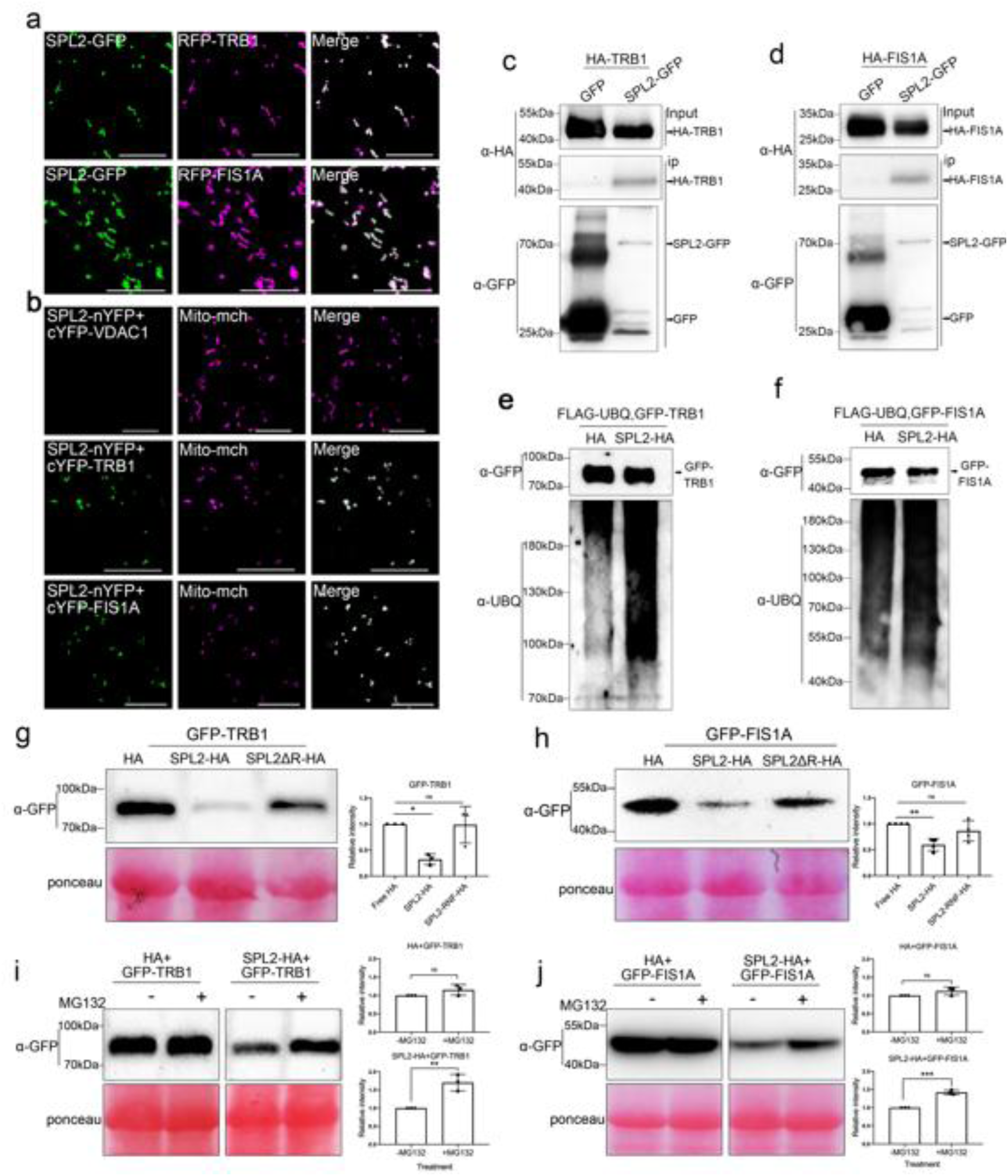
SPL2 interacts with TRB1 and FIS1A to promote their ubiquitin-mediated degradation. **a.** SPL2-GFP co-localises with GFP-TRB1 or GFP-FIS1A at the outer mitochondrial membrane in *N. benthamiana* leaf epidermal cells. Scale bars, 10 μm. **b.** BiFC assays indicate that SPL2 interacts with both TRB1 and FIS1A in *N. benthamiana* leaf epidermal cells, with VDAC1 included as a negative control. Scale bars, 10 μm. **c-d.** Co-IP analysis indicates that SPL2 interacts with both TRB1 and FIS1A using a GFP-Trap assay. HA-TRB1 (c) and HA-FIS1A (d) are pulled down only in the presence of SPL2-GFP (right) but not with free GFP (left). **e-f.** TRB1 and FIS1A are highly ubiquitinated in the presence of SPL2. At 48 h post-infiltration, GFP-TRB1 or GFP-FIS1A was purified and detected by immunoblotting using an anti-GFP antibody and an anti-UBQ (ubiquitin) antibody. **g-h.** Protein degradation assays. Full-length SPL2 protein promoted degradation of GFP-TRB1 and GFP-FIS1A in *N. benthamiana* leaf epidermal cells, while the SPL2 protein without the E3 catalytic ring domain (SPL2ΔR) failed to do so. Ponceau S staining was used as the loading control. At least three independent experiments were performed with similar results. **i-j.** SPL2-modulated degradation of TRB1 (i) and FIS1A (j) is markedly slowed in the presence of the proteasome inhibitor MG132. Free HA was used as a negative control. Three biological replicates were used for quantification. Error bars represent SD. Statistical significance was determined by one-way ANOVA with Dunnett’s test (g, h) or by a two-tailed Student’s *t*-test (i, j). Asterisks indicate statistically significant differences (\**P* < 0.05; \*\**P* < 0.01; \*\*\**P* < 0.001).

Since SPL2 bears E3 ligase activity^20^, we suspect it may ubiquitinate its interacting proteins. Using the *E. coli* reconstitution system^26^, we found that TRB1 ubiquitination occurred only in the presence of SPL2. Furthermore, deletion of the RING domain (SPL2ΔR) abolished this ubiquitination, confirming that SPL2-mediated ubiquitination of TRB1 depends on a functional RING domain (Extended Data Fig. 4a). Further tests using *N. benthamiana* confirmed that the expression of SPL2 is sufficient to induce TRB1 ubiquitination *in vivo* (Fig. 2e). The level of TRB1 decreased significantly in the presence of SPL2, and this effect could be restored after treatment with MG132, a proteasome degradation inhibitor (Fig. 2g, i). The experiments were repeated using FIS1A, yielding similar results, which suggest that SPL2 modulates mitochondrial protein degradation through ubiquitination (Fig. 2f, h, j). Furthermore, mass spectrometry analysis of TRB1 identified six lysine residues as potential ubiquitination sites; we mutated them to arginine. The TRB1^6KR^ mutation is still localised to mitochondria and interacts with SPL2, but it cannot be degraded (Extended Data Fig. 4b-k).

TRB1 is known to regulate ER-mitochondrial interaction at the ER-mitochondrial contact sites through the interaction with the ER-localised VAP27-1 protein^9^. We hypothesise that SPL2 and its interaction proteins also participate in this process. Y2H, co-IP, and BiFC assays collectively demonstrate FIS1A-TRB1-VAP27-1 form a complex (Extended Data Fig. 3b-c, Extended Data Fig. 5a-e). In VAP27-1-CFP, GFP-FIS1A and RFP-TRB1 triple-expressing cells, most of their co-localisation signal overlaps with the mitochondria and the closely associated ER membrane (Fig. 3a). When their BiFC constructs were co-expressed with the ER and mitochondrial marker, we found the interaction between FIS1A and VAP27-1 likely occurs at the ER-mitochondrial interface (Fig. 3b), like the interaction pattern between TRB1 and VAP27-1 (Extended Data Fig. 5f)^9^. Interestingly, some mitochondrial-localised SPL2 is closely associated with the ER membrane (Extended Data Fig. 5g) and interacts with VAP27-1, as indicated by the Y2H (Extended Data Fig. 3d) and BiFC studies (Extended Data Fig. 5h). Taken together, we proposed that FIS1A and SPL2 are likely part of the ER-mitochondrial tethering complex, in addition to VAP27-1 and TRB1.

**Fig 3.**
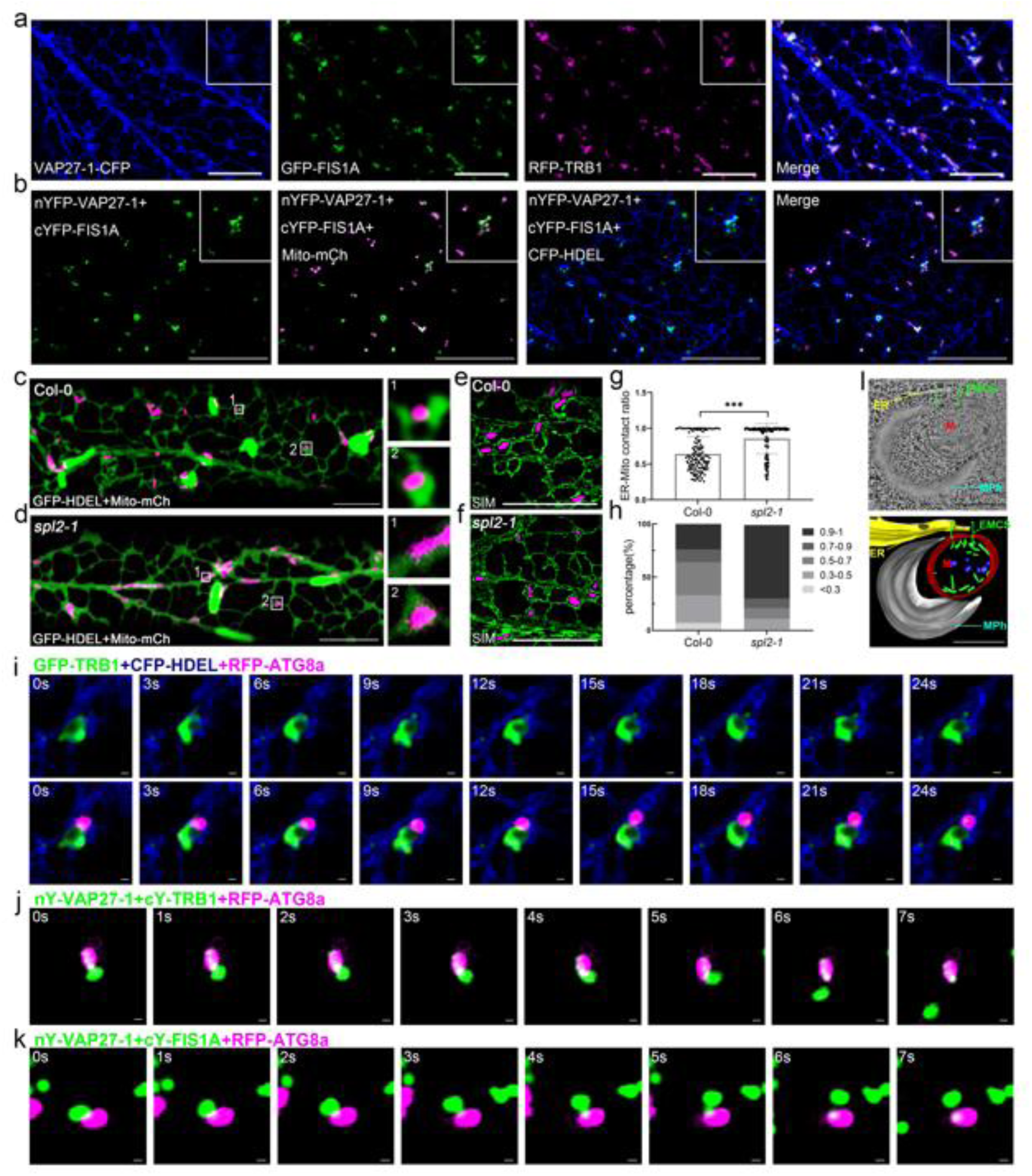
SPL2 modulates ER-mitochondria contacts (EMCSs) and mitophagosome formation through the TRB1-FIS1A-VAP27-1 complex. **a.** Triple expression of VAP27-1-CFP (blue), GFP-FIS1A (green), and RFP-TRB1 (magenta), shows partial co-localisation at ER-mitochondria contact sites in epidermal cells of *N. benthamiana* leaves. Scale bar, 10 μm. **b.** Split-YFP-based BiFC assays revealed that co-expression of nYFP-VAP27-1 and cYFP-FIS1A generated reconstituted YFP signals. These BiFC signals colocalised with both Mito-mCherry and CFP-HDEL, demonstrating that the VAP27-1-FIS1A interaction occurs at ER-mitochondria contact sites. Scale bar, 10 μm. **c-h.** Increased ER-mito contact ratio in etiolated hypocotyl cells of the *spl2* mutant relative to Col-0. **c-d.** Representative images showing ER (GFP-HDEL, green) and mitochondria (Mito-mCherry, magenta) in four-day-old hypocotyl cells of Col-0 and *spl2* stable lines. Enlarged views demonstrate the ER-mitochondria spatial relationships. **e-f**. Studtying ER-Mitochondria association in *Col-0* and *spl2* mutant by Super-resolution microscopy (SIM). **g.** Quantification of ER-mitochondrial association relative to total mitochondrial perimeter imaged as in (c, d). **h.** The frequency distribution of the ER-Mitochondria contact ratio. Data from 10 cells (n = 10); ≥130 mitochondria were used for the quantification. Scale bar, 10 μm. Error bars represent SD, and asterisks represent means with a significant difference at *P* < 0.05 (two-tailed Student’s *t*-test). **i.** Time-lapse imaging of CFP-HDEL (blue), GFP-TRB1 (green), and RFP-ATG8a (magenta) in *N. benthamiana*. The GFP-TRB1-labelled mitochondrial fragment forms at the ER-mitochondrial interface and is engulfed by RFP-ATG8a-labelled autophagosomes. Scale bar, 1 μm. **j-k.** Time-lapse imaging reveals that VAP27-1-TRB1- or VAP27-1-FIS1A-labelled ER-mitochondria contacts partially colocalise with ATG8a. nYFP-VAP27-1 and cYFP-TRB1 (j) or nYFP-VAP27-1 and cYFP-FIS1A (k) were co-expressed in *N. benthamiana.* Scale bar, 500 nm. **l.** Electron tomograms show mitochondria (M) in seven-day-old Arabidopsis roots after mitophagy induction (DNP treatment), with mitophagosomes (MPh) forming adjacent to ER-mitochondria contact sites. Scale bar, 200 nm.

As TRB1 positively mediates ER-mitochondrial interaction^9^, we suspected that SPL2 may modulate this interaction by controlling the levels of ER-mitochondrial tethering proteins. To test this hypothesis, we generated Arabidopsis lines expressing ER and mitochondrial markers and found a higher frequency of ER-mitochondrial association in the *spl2* mutant (Fig. 3c-f). The interaction between the ER and mitochondria is also calculated. The results indicate that mitochondria in the *spl2* mutant exhibit a higher ER coverage (ER-Mito contact ratio; Fig. 3g-h). The average distance between ER and mitochondria was quantified at the ultrastructural level with the same conclusion (Extended Data Fig. 6a-c). In cells with mitochondrial aggregation, aberrant ER aggregates and enhanced ER-mitochondria associations were also found (Extended Data Fig. 6d). In agreement with this observation, the protein level of endogenous TRB1 increased significantly in the *spl2* mutant, a change that may be attributed to the observed reduction in protein ubiquitination (Extended Data Fig. 6e-f).

In animals, the ER-mitochondrial contact sites are pivotal for the exchange of signalling molecules and lipids, essential for mitochondrial respiration and homeostasis. Furthermore, specific autophagy regulators are recruited to the ER-mitochondrial junctions, thereby enhancing membrane tethering and autophagosome formation^27–29^. To understand how TRB1 and FIS1A coordinate ER-mitochondrial interaction and mitophagy, time-lapse microscopy was performed to investigate their dynamic relationship. Over time, mitochondrial membrane fragments labelled with GFP-TRB1 are seen to be detach from the mitochondria at the ER-mitochondrial interface and concurrently engulfed by RFP-ATG8a labelled autophagosomes (Fig. 3i). Similar events were frequently found in cells expressing the BiFC constructs of VAP27-1, TRB1 and FIS1A: the VAP27-1-TRB1/FIS1A labelled EMCS (as indicated by the BiFC signal) partially co-localised with ATG8a, followed by mitochondrial fission and mitophagosome formation (Fig. 3j-k, Extended Data Fig. 7). We hypothesise that TRB1/FIS1A may facilitate the cleave of mitochondrial fragments that are engulfed by autophagy. The contact between ER, mitochondria and mitophagosomes is further confirmed by TEM tomography (Fig. 3l), using Arabidopsis root cells treated with 2,4-dinitrophenol (DNP), which is known to induce mitophagy (Extended Data Fig. 8a-b)^9,30^. Therefore, we believe that the EMCS-localised TRB1 and FIS1A are very likely key regulators of mitophagy, in coordination with their activities in ER-mitochondrial interaction.

In our previous study, we showed that TRB1 positively mediates mitophagy^9^. Here, we reconfirmed this result and show that TRB1 overexpression promotes mitochondrial turnover (as evident by the number of vacuole-accumulated mitochondria after Concanamycin A (Conc A) treatment), and its loss-of-function mutant (*trb1trb2*) shows an opposite effect (Extended Data Fig. 8c-e). Notably, we cannot exclude that micro-autophagy is also active and contributes to mitochondrial internalisation^31,32^. However, according to previous studies^9,10^, the number of mitochondria accumulated in vacuoles is sufficient to give a consistent estimate of the total mitophagy activity.

Since TRB1 levels are higher in the *spl2* mutant (Extended Data Fig. 6f), we suspect it is associated with abnormal mitophagy activity. As expected, more mitochondria are found in the vacuole of the *spl2* mutant after Conc A treatment, while the mitochondrial turnover is inhibited in SPL2 over-expressing lines (Fig. 4a-c). The activity of mitophagy was further confirmed biochemically using a transgenic Arabidopsis line expressing mitochondrial localized IDH1-GFP (mitochondrial isocitrate dehydrogenase 1; Extended Data Fig. 8f). It has been shown that IDH1-GFP degradation increased with high mitophagy flux^10^. We similarly observed that DNP treatment induced degradation of IDH1-GFP, resulting in the cleavage of GFP as a breakdown product (Extended Data Fig. 8g). In the *spl2* mutant, increased turnover of IDH1-GFP was evident as observed by increased levels of cleaved GFP (Fig. 4d-e), suggesting increased mitophagy activity.

**Fig 4.**
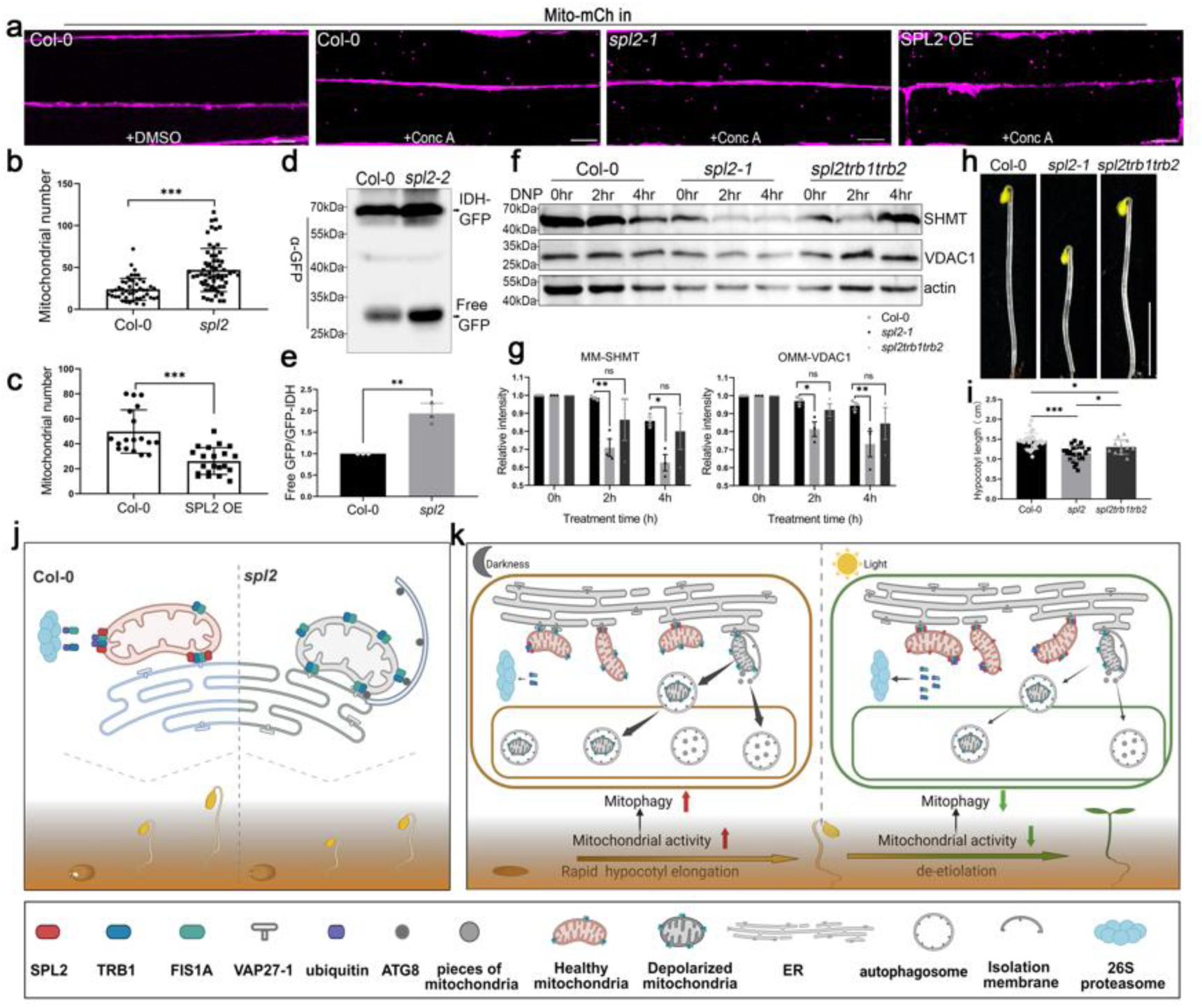
SPL2-mediated mitophagy is TRB1-dependent. **a.** Representative images of 5-day-old Arabidopsis etiolated hypocotyl cells expressing Mito-mCherry in Col-0, *spl2-1* mutant, and SPL2 OE lines after Conc A treatment, with little vacuolar signal observed in DMSO-treated control cells. Scale bar, 10 μm. **b-c.** Quantification of vacuolar mitochondria per 30×30 μm field in *spl2-1* mutants (b) and SPL2-overexpressing lines (c) after Conc A treatment. The results indicate that mitochondrial degradation is increased in *spl2* mutants compared to Col-0, whereas it is suppressed in SPL2-overexpressing lines. n ≥20 from 3 seedlings. Error bars represent SD. Asterisks indicate statistically significant differences from wild type (two-tailed Student’s *t*-test; *** *P* < 0.001). **d-e.** Mitophagy activity was significantly enhanced in *spl2-2* mutants, as determined using a transgenic Arabidopsis line expressing IDH1-GFP. 7-day-old Col-0 and *spl2-2* etiolated seedlings were used with three biological replicates for quantification. Error bars represent SD. Asterisks indicate statistically significant differences from wild type (two-tailed Student’s *t*-test; ** *P* < 0.01). **f.** Immunoblot of mitochondria matrix (MM) protein SHMT and outer mitochondrial membrane (OMM) proteins VDAC1 in Col-0 and *spl2-1* mutant seedlings treated with 50 μM DNP to induce mitophagy (0-4 h). Equal loading was verified by actin immunodetection. Protein levels were quantified relative to actin (loading control), with the level at 0 hours set to 1 for each genotype (Col-0, *spl2*, and *spl2 trb1 trb2*). **g.** Quantification of the immunoblot in (f) shows increased levels of SHMT and VDAC1 degradation in the *spl2* mutants, whereas mitophagy is suppressed in *spl2trb1trb2* triple mutants. Three biological repeats were used. Error bars represent SEM, and the asterisks represent means that are significantly different at *P* < 0.05 from a two-way ANOVA with Dunnett’s multiple comparisons test. **h-i.** The hypocotyl elongation defect of *spl2-1* mutants was partially restored in *spl2trb1trb2* mutants. Seedlings were grown in darkness for 4 days on 1/2 MS medium (Error bars represent SD; n=30). Asterisks indicate significantly different means at *P* < 0.05, as determined by a one-way ANOVA with Dunnett’s multiple comparisons test. Scale bar, 0.5 cm. **j.** Schematic illustrating that SPL2 loss-of-function disrupts seedling emergence from soil by enhancing ER-mitochondrial contact and mitophagy (through TRB1) compared to Col-0. **k.** Working model of the coordinated pathways between mitophagy and seedling emergence. In darkness, rapid hypocotyl elongation is sustained by high mitochondrial metabolic activity and increased mitophagy flux. The level of SPL2 is lower in this condition, allowing more TRB1 accumulation to promote mitophagy. Upon light exposure, the SPL2 protein accumulates, reducing mitophagy.

Next, we measured the levels of endogenous mitochondrial components as indicators of mitochondrial turnover, including the mitochondrial matrix protein, HMT (serine transhydroxymethyltransferase) and the outer mitochondrial membrane protein, VDAC (voltage-dependent anion channel)^30^. The levels of these two proteins were significantly reduced in the *spl2* mutant after mitophagy induction with DNP (Fig. 4f). In agreement with this result, the number of cytoplasmic mitochondria in the *spl2* mutant is also reduced (Fig. 1h-i). We then created a *spl2trb1trb2* triple knock-out mutant and found that the increased mitochondrial degradation of the *spl2* mutant is partially restored to the wild-type level in the triple knock-out mutants (Fig. 4f). Interestingly, the hypocotyl elongation defect of the *spl2* mutant is also partially restored in the *spl2trb1trb2* mutant (Fig. 4h-i), suggesting the SPL2-mediated mitophagy during hypocotyl elongation is dependent on TRB1. Taken together, our results suggest that SPL2 may function as a molecular switch regulating mitochondrial degradation. High SPL2 levels induce TRB1/FIS1A degradation and block mitophagy, whereas knockout of SPL2 expression leads to increased TRB1/FIS1A levels, which in turn cause excessive mitophagy, perturbing mitochondrial homeostasis and function.

SPL2 activity is likely be precisely controlled to maintain the balance between mitochondrial degradation and biogenesis. We suspect that SPL2 expression may change during specific stresses or developmental stages that induce damaged mitochondria. Therefore, the expression of SPL2 was examined under various growth conditions, and its transcript level is significantly lower in dark-grown conditions (Extended Data Fig. 9a-b). This result is consistent with a previous report indicating that mitophagy in plants is mainly associated with dark starvation ^10^. The upregulation of SPL2 protein after light perception was further confirmed using a *spl2* complemented line (pSPL2::SPL2-GFP/*spl2-1*, Extended Data Fig. 9c). Such a difference may be related to seedling emergence, where seedlings initially exhibit rapid hypocotyl elongation in the dark, which is more likely to require higher mitochondrial output and mitophagy activity. However, hypocotyl elongation stops after light signal perception, so the demand for mitophagy is reduced (Extended Data Fig. 9d-i). Consequently, SPL2 expression increases to remove mitophagy regulators (Extended Data Fig. 9c).

To assess the conservation of SPL2 function in seedling emergence and mitochondrial functional regulation among seed plants, we examined tomato and citrus orthologs and found that both SlSPL2-GFP and CsSPL2-GFP localise to mitochondria (Extended Data Fig. 10a). Further evidence was obtained using the *slspl2* CRISPR mutants, whose mitophagy activity, hypocotyl development, and seedling emergence are also significantly affected (Extended Data Fig. 10b-k).

In conclusion, we identify SPL2, a mitochondrial E3 ligase, as essential for maintaining energy homeostasis, promoting hypocotyl elongation, and facilitating seedling emergence. SPL2 interacts with mitochondrial outer membrane proteins (TRB1, FIS1A) and ER membrane proteins (VAP27-1), forming a complex at the EMCS. SPL2 acts as an upstream inhibitor of mitophagy, through mediating the ubiquitin-dependent degradation of the mitophagy receptors and ER-mitochondrial tethers. However, it is not clear how SPL2 recognises damaged mitochondria; and it could be speculated that it may act in concert with other mitophagy regulators, such as ELM1 and DRP3, which mediate mitochondrial fission and mitophagosome formation^33^. During seedling emergence and light perception, SPL2 expression negatively correlates with mitophagy, and hypocotyl elongation, suggesting that SPL2 may contribute to photomorphogenesis through an as-yet uncharacterized mechanism. The underlying mechanism could be an intriguing question for future studies. Our work here provides evidence that ubiquitination pathways are indeed involved in regulating mitochondrial quality control in plants and offers mechanistic insights into seedling emergence from an organelle-interaction perspective.

## Acknowledgements

We thank Prof. Ronghui Pan (Zhejiang University), Shangwei Zhong (Peking University), and Prof. Qihua Ling (CEMPS) for their advice throughout the project. We thank Patrick Hussey (Durham University) for his advice and long-term support. We thank Prof Peter Pimpl (SUSTech) for providing the construct of NbG. We thank the Public Laboratory of Electron Microscopy at Huazhong Agricultural University, the Center for Protein Research (CPR) at Huazhong Agricultural University, the Core Facilities of the College of Life Science and Technology, and the Microscopy Platform of the Key Laboratory of Horticulture for their assistance; Guangzhou CSR Biotech Co., Ltd. for live-cell imaging using its commercial super-resolution microscope (HIS-SIM); and Airy Technologies Co., Ltd. (China) for live-cell fluorescence imaging utilizing its Polar-SIM microscope (Nova-SD). The project was supported by the National Natural Science Foundation of China (NSFC) grants (32261160371, 92254307), the Science and Technology Project of Hubei Province (2024AFE005), the Fundamental and Interdisciplinary Disciplines Breakthrough Plan of the Ministry of Education of China (JYB2025XDXM701), the Fundamental Research Funds for the Central Universities (2662023PY011), the Young Scientist Fostering Fund of the National Key Laboratory for Germplasm Innovation and Utilization of Horticultural Crops (Horti-PY-2023-001), and the Research Grant Council (RGC) of Hong Kong grants (GRF14113921, GRF14109222, GRF14110823, GRF14113424, N_CUHK462/22, C4014-23G) awarded to B.-H.K.

## Author Contributions

Z.T. and Y.H. performed most of the experiments and wrote the manuscript with P.W.; P.W. designed the experiment and supervised the project; P.D. edited the manuscript; C.L. and P.D. were involved in the early mitophagy studies and generated various TRB1-related materials; Q.Z. performed the TEM studies; F.H. helped with plant transformation and phenotype studies; J.M and J.L performed the tomography studies B.K., X.Q. and Y.C. participated in project discussions and result interpretation throughout the work.

## Online Methods

### Generating fluorescent protein constructs

To generate fluorescent protein fusion constructs of SPL2, the coding sequences (CDSs) of various SPL2 homologs (Arabidopsis, tomato and Citrus) were amplified from cDNA libraries of Arabidopsis, tomato, and citrus using Phanta Max Super-Fidelity DNA Polymerase (Vazyme, P505). AtSPL2 was cloned into the pRI121 vector (driven by the 35S promoter) with C-terminal GFP or RFP tags via Gibson assembly, whereas SlSPL2 and CsSPL2 were cloned into pMDC83-GFP (containing a C-terminal GFP tag). The coding sequences of Arabidopsis TRB1 and FIS1A were amplified in the same way and cloned into pMDC43-GFP (N-terminal GFP), pK7WR2 (N-terminal RFP), or pK7WHA (N-terminal HA) vectors using Gateway cloning (Invitrogen). For the expression of SPL2 under its native promoter, approximately 1500 bp of the promoter region directly upstream of the SPL2 start codon, and its genomic sequence were cloned into pCAMBIA1301 to generate pSPL2::SPL2-GFP via Gibson assembly using the HindIII and BamHI restriction sites. To produce a mitochondrial matrix marker for the mitophagy flux test, the full coding region of IDH1 (mitochondrial isocitrate dehydrogenase) was amplified from an Arabidopsis cDNA library and cloned into pMDC83-GFP via Gateway cloning, generating the IDH1-GFP construct. To generate the mRFP-NbG construct, the sequence was amplified from the expression construct provided by Prof. Peter Pimpl^34^. The mRFP-NbG fragment was cloned into the pRI121 vector between the XbaI and SacI restriction sites using Gibson cloning to create the pRI121-mRFP-FLAG-NbG construct. Other constructs used in this study were published previously; please refer to extended Data Table 1 for details. All primers used in this study are shown in extended Data Table 2.

### Plant transformation and growth conditions

Arabidopsis seeds were surface sterilized with 70% (v:v) ethanol for 10 min, washed 6 times with sterile water, then sown on half-strength Murashige and Skoog (MS) medium (Duchefa) supplemented with 1% (w:v) sucrose, 0.05% (w:v) 4-morpholineethanesulfonic acid (MES) and 0.6% (w:v) agar at pH 5.8-6.0. Seeds were imbibed for 2 d at 4°C in darkness, and seedlings were grown under the photoperiodic cycle of 16 h light at 22°C and 8 h dark at 18°C. Stable transgenic Arabidopsis lines expressing Mito-mCherry (mitochondria), GFP-HDEL (ER), CFP-PT (plastids), and GFP-ATG8a (autophagosomes) have been described previously^35–37^. To generate Arabidopsis lines stably expressing IDH1-GFP and SPL2-GFP, wild-type Arabidopsis (Col-0) was transformed via Agrobacterium-mediated floral dipping^38^. The Mito-mCherry stable line was transformed with pSPL2::SPL2-GFP, 35S::SPL2-GFP or 35S::GFP-TRB1 via floral dipping, generating the respective double-expressing lines. Arabidopsis co-expressing 35S::SPL2-GFP + Mito-mCherry + CFP-PT or pSPL2::SPL2-GFP + Mito-mCherry + CFP-PT were generated by crossing the CFP-PT single-expressing line with the SPL2 + Mito-mCherry double-expressing lines. Please refer to extended Data Table 3 for all transgenic lines used in this study.

The *trb1trb2* double knockout mutants used in this study were previously published^9^. The *spl2* single mutants were generated using the CRISPR-Cas9 system, and two independent lines (*spl2-1 and spl2-2)* were selected for phenotypic studies with similar results. Therefore, the *spl2-1* line was used for further cell biological and biochemical analysis to avoid replications. For the SlSPL2 CRISPR/Cas9 constructs, the target sequences were designed using the online CRISPR-P 2.0 tool (http://crispr.hzau.edu.cn/CRISPR2/)^39^, and two single-guide RNA (sgRNA) expression cassettes were assembled into the pTX vector via Gibson assembly using BsaI restriction sites. The *spl2 trb1 trb2* triple mutants were generated by genetic crossing. The IDH1-GFP/*spl2* and Mito-mCherry/*spl2* transgenic lines were generated by crossing stable transgenic plants expressing IDH1-GFP or Mito-mCherry with *spl2* mutants. Please refer to extended Data Table 3 for all transgenic lines used in this study.

The tomato (*S*. *lycopersicum*) cultivar MicroTom was used for genetic transformation, generating the *slspl2* CRISPR mutant. To do this, tomato seeds were surface-sterilised by immersion in 70% (v/v) ethanol for 1 min, followed by two washes with sterile water. Seeds were then treated with 7% NaClO for 15 min and rinsed four times with sterile water. After drying on sterile filter paper, seeds were germinated on 1/2 MS medium. Tomato transformation was mediated by Agrobacterium tumefaciens (strain GV3101) using the leaf disc method, as described in previous studies^40^. The GFP-TOM20-3/*slspl2* transgenic lines were generated by crossing stable transgenic plants expressing GFP-TOM20-3.

### Soil emergence assay

Soil emergence assays were performed as previously described^41^. In brief, seeds were surface-sterilised and plated on 1/2 MS medium in 50 cm² plates. After storage at 4°C in the dark for 3 days, the plates were illuminated with white light for 6 hours. Next, 10 ml of 40-80 mesh sand (autoclaved, Sigma) was spread evenly over the medium, covering the seeds to a depth of 3 mm. The bottom and rim of each plate were covered with aluminium foil to prevent root exposure to light. The plate top was then illuminated with white light for the indicated number of days.

### *N. benthamiana* transient expression

*N. benthamiana* plants were grown with 16 h of light (25 °C) and 8 h of darkness (18 °C). Agrobacterium carrying different constructs was cultured in Luria-Bertani liquid medium at 28 °C overnight. And the agrobacteria were resuspended in infiltration buffer containing 100 μM acetosyringone, 50 mM MES, and 2 mM Na3PO4. Leaves of four-week-old *N. benthamiana* plants were used for transient transformation through Agrobacterium infiltration, as described in Sparkes et al., 2006^42^. Optimal expression of fluorescent fusion protein is achieved using an OD600 at 0.05-0.2, and the transformed leaf segments were studied 48-60 hours after infiltration.

### Co-IP assay

For co-immunoprecipitation, 0.2 g of plant material was ground in liquid nitrogen, and proteins were extracted by adding Nonidet P-40 buffer containing 1 mM phenylmethylsulfonyl fluoride (PMSF). The mixture was incubated on ice for 30 min, then centrifuged at 12,000 × g for 10 min at 4 °C three times. The supernatant was incubated with 10 μl of GFP-Trap Agarose beads (Chromotek, gtma-20) for 1 h at 4 °C, followed by centrifugation at 2,500 × g for 2 min at 4 °C. The beads were washed three times with wash buffer (10 mM Tris-HCl, pH 7.5, 150 mM NaCl, 0.5 mM EDTA, 1 mM PMSF, 0.5% Triton-X100, and Complete Protease Inhibitor [Sigma]). Finally, the beads were eluted in 50 μl of 2× SDS loading buffer, boiled at 95 °C for 10 min, and the agarose pellet was collected for western blot analysis.

### *In vitro* protein ubiquitination assay

Ubiquitination assay in *E. coli* was performed as previously described^26^. Either pACYCDuet-Myc-UBC8-S, pACYCDuet-SPL2-Myc-UBC8-S, or pACYCDuet-SPL2ΔRNF-Myc-UBC8-S, together with pCDFDuet and pET-28a-FLAG-UBQ, were mixed in equal amounts and co-transformed into *E. coli* BL21(DE3) competent cells via a standard protocol. *E. coli* BL21 (DE3) strains containing different plasmid combinations were cultured in LB medium at 37°C. When the OD600 reached 0.4–0.6, protein expression was induced by adding 0.5 mM IPTG. After induction, the cultures were grown at 28°C for 10–12 h and then stored at 4°C overnight. Bacterial cells from 500 µL of culture were harvested by centrifugation at 12,000 × g for 5 min. The pellet was resuspended in 100 µL of 1× SDS-PAGE sample buffer and boiled at 95°C for 5 min. The crude extracts were then separated by SDS-PAGE and analysed by immunoblotting with corresponding antibodies.

### Protein ubiquitination and degradation assay *in vivo*

To investigate SPL2-mediated protein ubiquitination, the SPL2-HA construct was co-expressed with target proteins (e.g., TRB1 or FIS1A) in *N. benthamiana* leaves via Agrobacterium-mediated infiltration. Approximately 0.2 g of leaf tissue was cryoground in liquid nitrogen, and total proteins were extracted using NP-40 buffer supplemented with 1 mM PMSF. GFP-tagged target proteins (GFP-TRB1 or GFP-FIS1A) were immunoprecipitated with anti-GFP beads (Chromotek, gtma-20), followed by immunoblotting with anti-HA antibody (1:5000, Yeasen) to detect ubiquitinated species and anti-Ub antibody (1:5000, Agrisera) to confirm polyubiquitination. For protein degradation assays, total proteins from *N. benthamiana* or transgenic Arabidopsis were extracted in 2× SDS buffer. GFP fusion proteins were detected by immunoblotting using an anti-GFP antibody (1:5000, BBI). As a control, transformed leaf sections were pre-treated with the 26S proteasome inhibitor MG132 (50 μM; Sigma) 4 hours before protein extraction. Additionally, 8-day-old Arabidopsis seedlings (Col-0 and *spl2* mutant) were analysed, with endogenous TRB1 levels detected using a validated anti-TRB1 antibody as previously described.

### LC-MS screen and ubiquitination sites identification

To identify the putative ubiquitination site of TRB1, total protein was extracted from 6-day-old Arabidopsis plants expressing 35S::GFP-TRB1, and the protein was immunoprecipitated using GFP-Trap beads. The immunoprecipitated complexes were separated by SDS-PAGE, and the gel region corresponding to the molecular weight of GFP-TRB1 was excised. Following in-gel tryptic digestion, the resulting peptides were analysed by mass spectrometry to identify ubiquitination sites. The mass spectrometry proteomics data have been deposited to the ProteomeXchange Consortium (https://proteomecentral.proteomexchange.org) via the iProX partner repository with the dataset identifier PXD068375. In addition, we performed an in-silico analysis of potential ubiquitination sites (http://ubpred.org/). Combining LC-MS screening and computational prediction, six putative sites were identified.

### Immunoblot analysis

Arabidopsis seedlings or *N. benthamiana* leaf segments were ground in liquid nitrogen and mixed with an equal volume of 2× SDS buffer (50 mM Tris-HCl, 40 mM NaCl, 10 mM MgCl₂, 5 mM EDTA, 5 mM DTT). The mixture was incubated at 95 °C for 10 min, then centrifuged at room temperature to collect the supernatant. An appropriate amount of total protein was loaded onto a 5% stacking gel and separated by 10% or 12% SDS-PAGE. Following electrophoresis, proteins were transferred to a nitrocellulose membrane. For detection, the membrane was blocked with 5% milk in 2× TBST buffer, then incubated with primary antibodies (anti-TRB1 (1:500), anti-VDAC1 (1:5000, Agrisera), anti-GFP (1:5000, BBI), anti-HA-Tag (1:5000, Yeasen), anti-Actin (1:1000, BBI) and horseradish peroxidase (HRP)-conjugated goat anti-mouse/rabbit secondary antibodies (Yeasen/ BBI, 1:5000) at room temperature. Original images of all immunoblots are provided in the Source data document.

### Autophagy assays

For mitophagy flux analysis, Arabidopsis Col-0 and *spl2* mutant seedlings stably expressing IDH1-GFP were generated; 0.2 g of plant tissue was ground in liquid nitrogen, and total proteins were extracted using 2× SDS buffer. Western blotting was performed using an anti-GFP antibody to assess mitophagy flux through the analysis of GFP cleavage products, which reflect autophagic degradation of mitochondria. For Conc A treatment, Arabidopsis (or tomato) seedlings were initially grown on 1/2 MS medium, then transferred to liquid MS medium or KCMS medium supplemented with Conc A (1 μM, Sigma) for 12 h. Fluorescence images were captured using a Leica SP8 confocal microscope, with excitation wavelengths adjusted according to the expressed fluorescent proteins. For mitochondrial protein degradation analysis, 8-day-old Arabidopsis seedlings were transferred to liquid MS medium and treated with DMSO (control) or DNP (50 μM, Sigma) for 0–4 h. Seedlings were harvested at each time point, flash-frozen in liquid nitrogen, and subjected to western blot analysis.

### BiFC analysis

For BiFC analysis, the full-length CDS of TRB1, FIS1A, and VDAC1 were cloned into pMDC43-cYFP plasmids (N-terminal cYFP fusion), and the full-length CDS of TRB1 and VAP27-1 were cloned into pMDC43-nYFP (N-terminal nYFP fusion). The full-length CDS of SPL2 was cloned into pEARLY-nYFP (C-terminal nYFP fusion). All constructs were transformed into the Agrobacterium tumefaciens strain GV3101 for plant transformation. Transiently transformed *N. benthamiana* leaves were imaged two days after infiltration using a Leica SP8 confocal microscope. For nYFP/cYFP detection, samples were excited at 514 nm and emissions detected at 550–580 nm. For multi-colour imaging (CFP/YFP/RFP), samples were excited at 405/514/552 nm and emissions detected at 450–480/550–580/590–640 nm, respectively.

### Yeast two-hybrid assays

Yeast two-hybrid experiments were performed using the split-ubiquitin dual membrane system^43^. The SPL2 coding sequences (without the stop codon) were fused to the Cub-LexA-VP16 fragment in the pBT3-STE vector via Gibson assembly using the SfiI restriction site. We selected sixteen proteins that are localized to the outer mitochondrial membrane and are known to regulate mitochondrial activities in various aspects, such as TOM7 and TOM40, core components of the TOM receptor complex responsible for recognizing and translocating cytosolically synthesized mitochondrial preproteins^44^; VDAC1, VDAC2, and VDAC3, which may facilitate metabolite exchange (also likely to regulate mitophagy)^45^; TRB1, MIRO1, and MIRO2, which regulate mitochondrial-ER interactions^9,21^; and DRP3a, FIS1A, FIS1B, PMD1, and PMD2, which are likely involved in mitochondrial fission^22,23,46^. Additionally, we also included TTM1 and DGS1 for their essential roles in mitochondrial function, and SP1 as a homolog of SPL2^47,48^. Full-length cDNAs of these mitochondrial outer membrane proteins were fused to the NubG fragment in the pPR3-N vector, which harbours a mutation in nUb to severely reduce its affinity for cUb.

To assess interactions within the TRB1-FIS1A-VAP27-1 complex, FIS1A and VAP27-1 were cloned into the pBT3-N vector. pPR3N-nUb and pPR3N-nUbG were used as positive and negative controls, respectively. Both prey and bait plasmids were co-transformed into the yeast NMY51 strain and cultured on SD/-Leu-Trp selective plates at 30 °C for 2–3 days. Co-transformed yeast clones were diluted (1:10 or 1:100) and spotted on selective media for growth assessment. 1 mM 3-AT was used to block leaky expression of the reporter gene.

### Transmission electron microscopy

For chemical fixation of Arabidopsis hypocotyls, 5-mm segments from 4-day-old Col-0 and *spl2* mutant seedlings were used. Samples were prefixed in 2.5% glutaraldehyde (v/v in 0.1 M phosphate buffer, pH 7.2) for 2 h, then rinsed three times with 0.1 M phosphate buffer (pH 7.2). Post-fixation was performed according to Li et al.,2022.^9^ Samples were embedded in SPI-PON 812 resin and polymerised at 60 °C for 48 h. Ultrathin sections (80 nm) were prepared using an EM UC7 Ultracut ultramicrotome (Leica, UC7), then observed and imaged with a transmission electron microscope (Hitachi H-7650) at an accelerating voltage of 80.0 kV. For mitochondrial morphology analysis, at least 50 mitochondria from three independent biological replicates were evaluated. For ER-mitochondria contact quantification, the shortest distance between mitochondria and adjacent ER membranes was measured using Fiji, with at least 50 mitochondria analysed from three independent biological replicates.

For electron tomography, TEM samples were prepared by high-pressure freezing followed by processing as previously described^49,50^. Briefly, seven-day-old Arabidopsis roots were incubated in 50 μM DNP and rapidly frozen using a high-pressure freezer. (Leica Microsystems, HPM100). An automatic freeze-substitution machine (Leica Microsystems, AFS2) was used to allow slow freeze-substitution with 2% OsO4 (Electron Microscopy Sciences, 19130) from −80 °C to room temperature. Excess OsO4 were washed away with acetone. Samples were then embedded in EPON resin (Electron Microscopy Sciences, 14120) by gradually increasing the resin concentration from 10%, 25%, 50%, 75%, to 100% diluted in acetone and polymerised at 60 °C overnight. EPON-embedded root samples were then cut into serial sections (250 nm) and collected on slot grids. After post-staining with uranyl acetate and lead citrate, a 200 kV Talos F200C electron microscope (Thermo Fisher, USA) was used to collect a double-axis tilt series from -60 ° to 60° at 1.5° increments. Dual-axis tomograms were reconstructed from pairs of image stacks with the etomo program of the IMOD software package (University of Colorado Boulder, USA) software package (http://bio3d.colorado.edu/imod/doc/3dmodguide.html) as previously described^51^.

### Light microscopy and image analysis

For subcellular localisation of SPL2, mitochondrial activity, and autophagosome analysis, images were collected using a Leica TCS SP8 laser-scanning confocal microscope with a 63× oil-immersion or 40× water-immersion objective lens. Multifluorescence imaging was performed in multi-track mode with line switching to avoid crosstalk. Time-lapse imaging was performed using an Olympus SpinSR10 spinning-disk confocal system equipped with a 63×/1.4 oil immersion objective. Super-resolution imaging analysis of SPL2 subcellular localisation was performed using the commercial Polar-SIM system (Airy Technologies Co., Ltd., China). Images were acquired with a 100×/1.49 NA oil immersion objective (Nikon, Japan). SIM image reconstruction was conducted using Airy-SIM software with dark-field pre-processing (dark correction). Super-resolution imaging of ER and mitochondria was performed using commercialized HIS-SIM (High Intelligent and Sensitive SIM) provided by Guangzhou CSR Biotech Co. Ltd. Images were acquired using a 100×/1.5 NA oil immersion objective (Olympus). The fluorescent intensity and band intensity (in the western blot assay) were measured using Fiji.

### Statistical analysis

Statistical analyses were performed using GraphPad Prism (v8.0). Data were analysed using appropriate statistical analysis methods, including Student’s *t*-tests and one-way ANOVA for multiple comparisons. A *P* value < 0.05 was considered statistically significant with asterisks denoting thresholds (* *P* < 0.05, ** *P* < 0.01, *** *P* < 0.001).

## Accession numbers

Accession numbers of genes in this article are AtSPL2 (AT1G54150), SlSPL2(Solyc12G049330), CsSPL2(Cs1G17520), SP1 (AT1G63900), TRB1 (AT1G05270), FIS1A (AT3G57090), FIS1B (AT5G12390), VAP27-1 (AT3G60600), MIRO1 (AT5G27540), MIRO2 (AT3G63150), PMD1 (AT3G58840), PMD2 (AT1G06530), DRP3A (AT4G33650), VDAC1 (AT3G01280), VDAC2 (AT5G67500), VDAC3 (AT5G15090), BAG1 (AT5G52060), TTM1 (AT1G73980) , DGS1 (AT5G12290) , TOM40 (AT3G20000) , TOM7(AT5G41685) , TOM20-3(AT3G27080) , IDH1 (AT4G35260)

## Extended data

**Extended Data Fig. 1.**
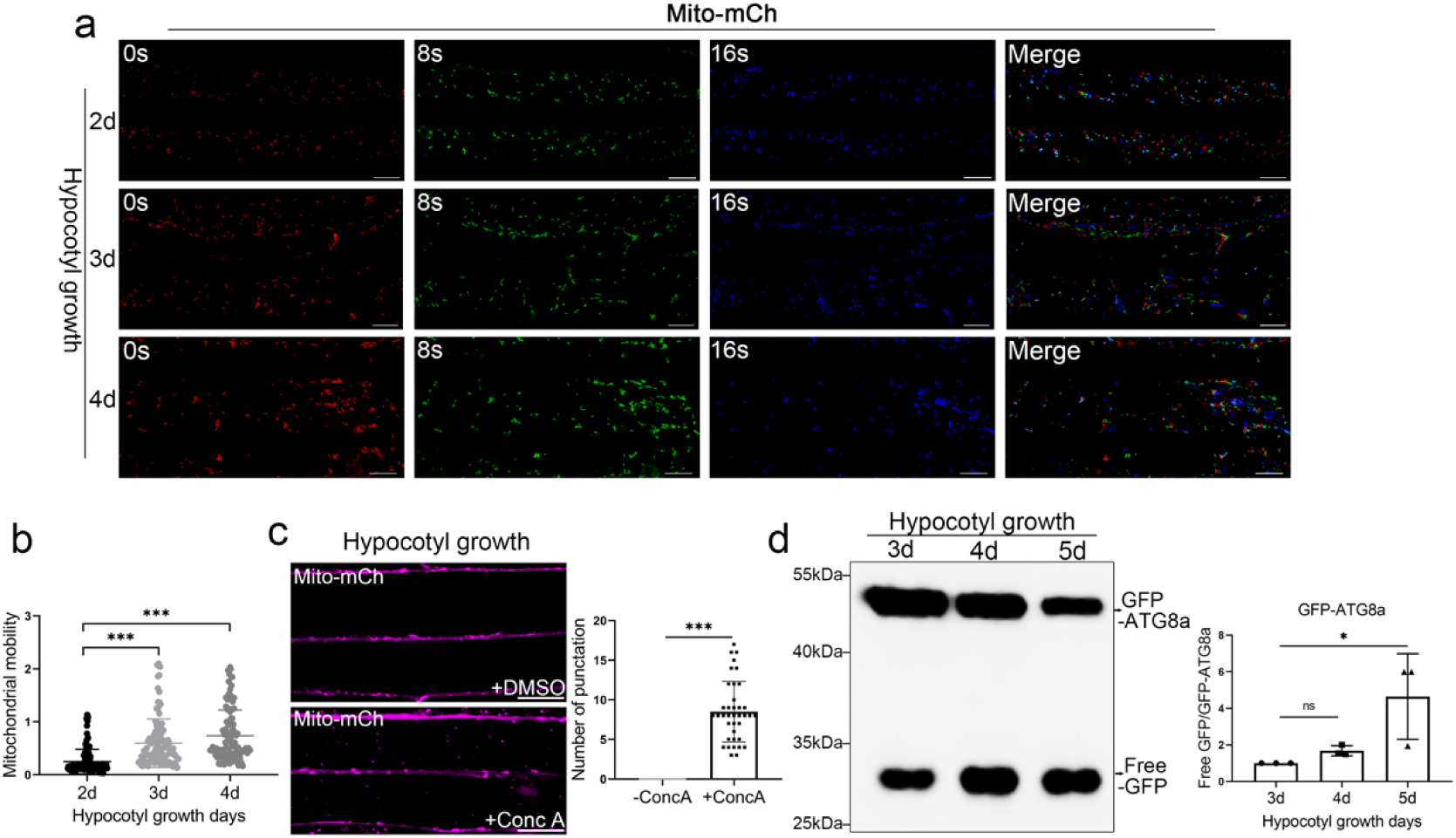
Mitochondrial dynamics, morphology and mitophagy flux analysis of hypocotyl during rapid elongation. **a-b.** Mitochondrial dynamics undergo profound changes in dark-grown hypocotyl cells. **a.** Time-lapse confocal imaging of mitochondrial dynamics in etiolated hypocotyl cells of Arabidopsis thaliana stably expressing Mito-mCherry. Scale bar, 10 µm. **b.** Quantification of mitochondrial motility in dark-grown hypocotyl cells. Error bars represent SD, and the asterisks represent means that are significantly different at *P* < 0.05 from a one-way ANOVA with Dunnett’s multiple comparisons test. **c.** The vacuolar accumulation of mitochondria (Mito-mCherry) in 4-day-old Arabidopsis seedlings after Conc A treatment. Scale bar, 10 μm. Error bars indicate SD, andat *P* < 0.05, as determined by a two-tailed Student’s *t*-test. **d.** The GFP-ATG8a cleavage assay reveals elevated autophagic flux in etiolated seedlings. Immunoblot analysis with anti-GFP antibody detects both full-length fusion proteins and free GFP. At least three independent experiments were performed with similar results. Error bars represent SD. Asterisks indicate significantly different means at *P* < 0.05, as determined by a one-way ANOVA with Dunnett’s multiple comparisons test.

**Extended Data Fig. 2.**
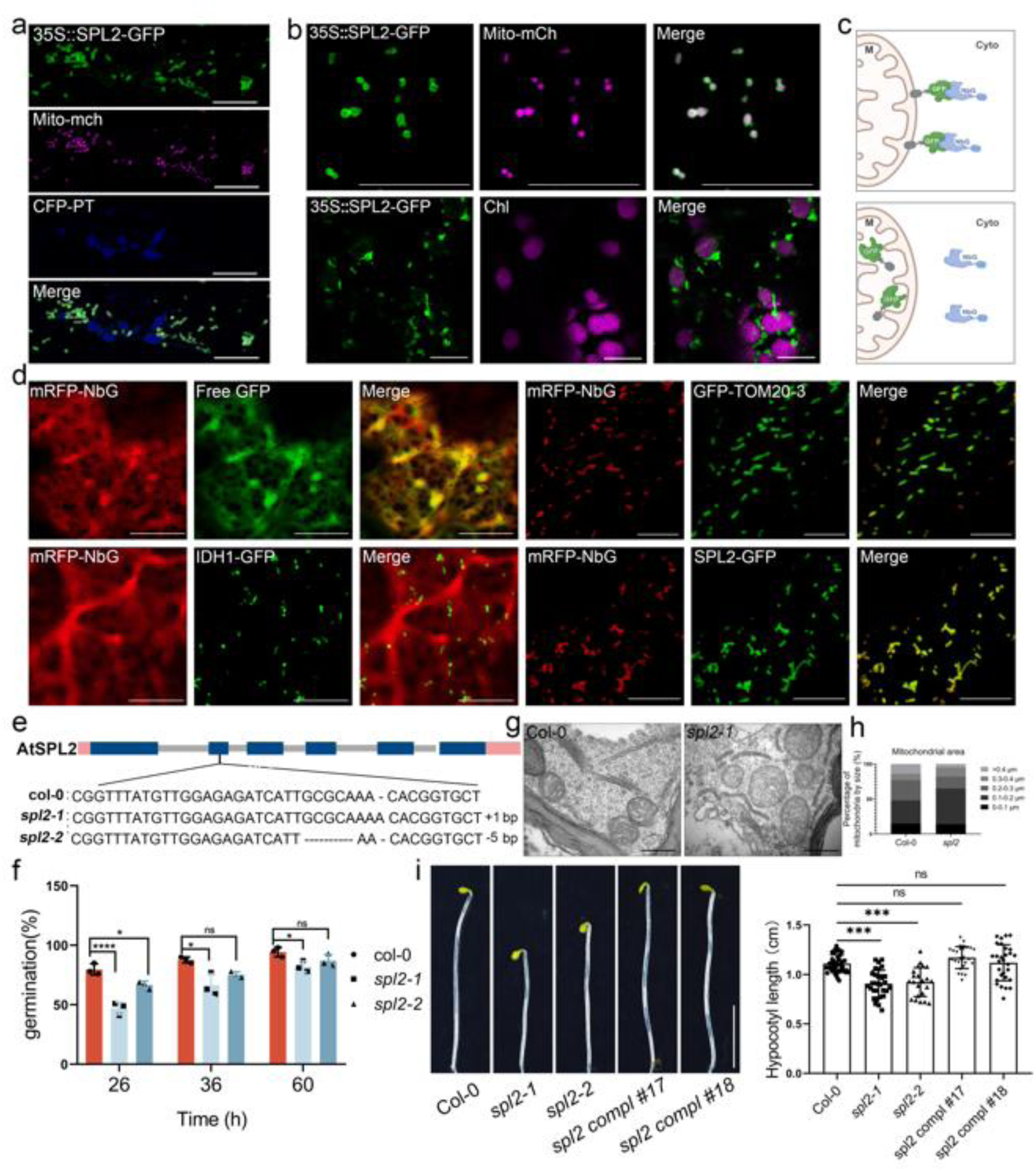
SPL2 is localised to the mitochondrial outer membrane and affects mitochondrial morphology and seed germination. **a.** In Arabidopsis hypocotyl cells, 35::SPL2-GFP predominantly localises to the mitochondrial outer membrane, as confirmed by the co-localisation with the matrix marker Mito-mCherry. Scale bar, 10 μm. **b.** Similarly, 35::SPL2-GFP predominantly localises to the mitochondrial outer membrane in *N. benthamiana* epidermal cells, as shown by co-expression with Mito-mCherry (mitochondria) with Chloroplast autofluorescence. Scale bar, 10 μm. **c.** Schematic illustrating the recruitment of mRFP-NbG to GFP-tagged proteins localised on the outer mitochondrial membrane (OMM). **d.** mRFP-NbG can be recruited to the mitochondria in the presence of GFP-TOM20-3 and SPL2-GFP (mitochondrial outer membrane), but not by IDH1-GFP (mitochondrial matrix) in *N. benthamiana* epidermal cells. Scale bar, 10 μm. **e.** Schematic diagram of the AtSPL2 gene and the CRISPR site. Two independent homozygous mutants, *spl2-1* and *spl2-2*, were selected for functional studies. **f.** In *spl2* mutants, seed germination is significantly altered. The germination time of Col-0, *spl2-1*, and *spl2-2* seeds were quantified at indicated time points post-transfer to standard growth conditions. **g-h.** TEM analysis of 3-day-old dark-grown seedlings revealed that mitochondrial fragmentation and aggregates were abundant in the *spl2* mutant. At least 50 mitochondria from 3 seedlings of at least 3 biologically independent samples were analysed. Scale bar, 500 nm. **i.** Gene complementation assay using the pSPL2::SPL2-GFP construct confirmed that the *spl2* mutant phenotype is due to SPL2 dysfunction. Scale bar, 0.5 cm. Error bars represent SD; asterisks indicate significant differences (*P* < 0.05, one-way ANOVA with Dunnett’s test; panels f and i).

**Extended Data Fig. 3.**
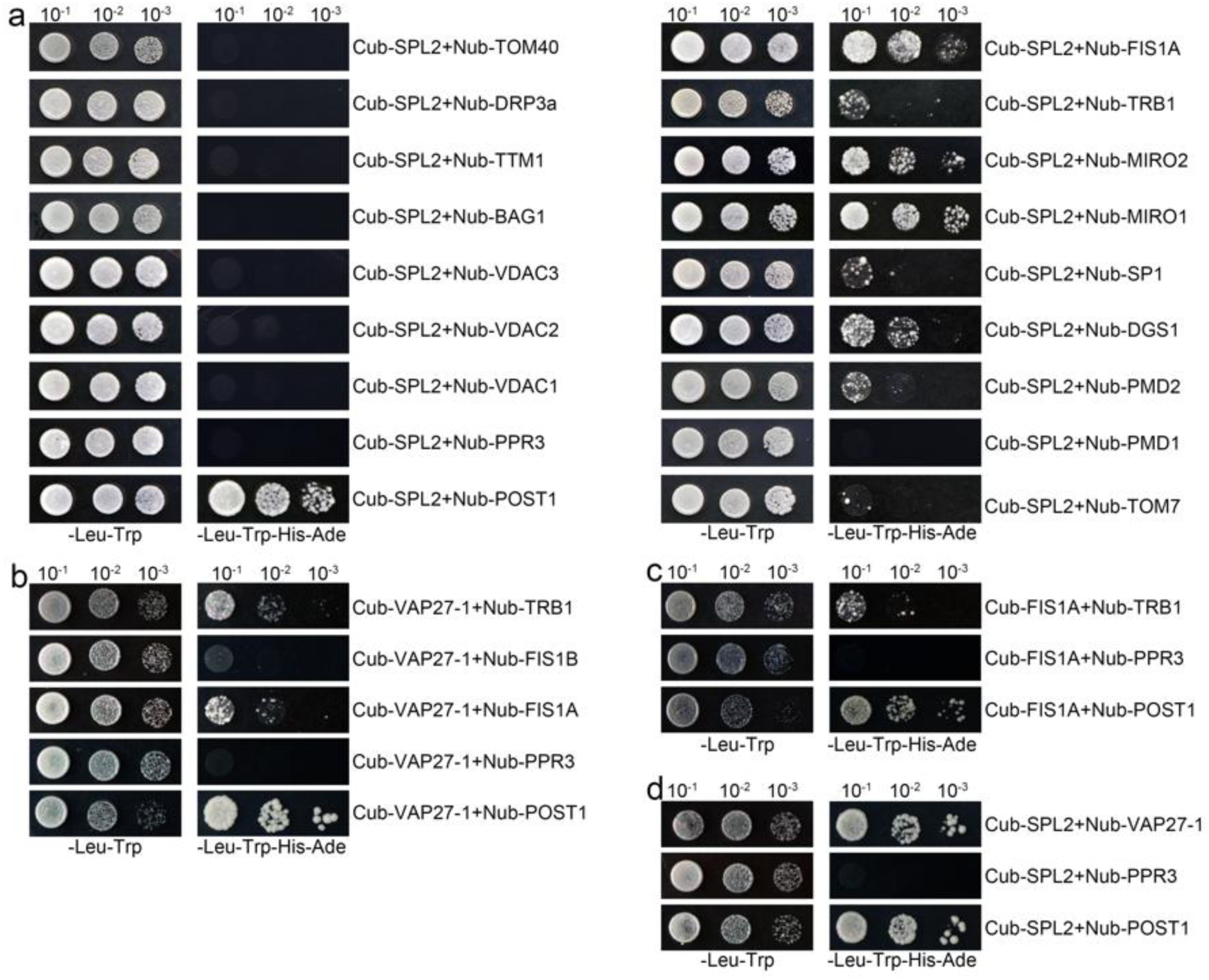
Yeast two-hybrid (Y2H) assays for putative protein interactions. **a.** SPL2 interacts with multiple mitochondrial outer membrane proteins in Y2H one-on-one assays. Sixteen candidate proteins were selected, and the detailed criteria are described in the Materials and Methods section under the subsection on yeast two-hybrid assays. **b.** Y2H assays show that VAP27-1 interacts with TRB1, FIS1A, and FIS1B. **c.** Y2H assays indicate a direct FIS1A-TRB1 interaction. **d.** Y2H assays show an interaction between SPL2 and VAP27-1. In all assays, NubG-PPR3 (negative control) and NubG-POST1 (positive control) were included. Yeast cells were used at an optical density at 600 nm (OD600) of 1, with 1:10 and 1:100 serial dilutions, on vector-selective media. All interactions were confirmed by growth assays on the selective media (-Leu/-Trp/-His/-Ade), and plates were incubated at 30°C for 3 days.

**Extended Data Fig. 4.**
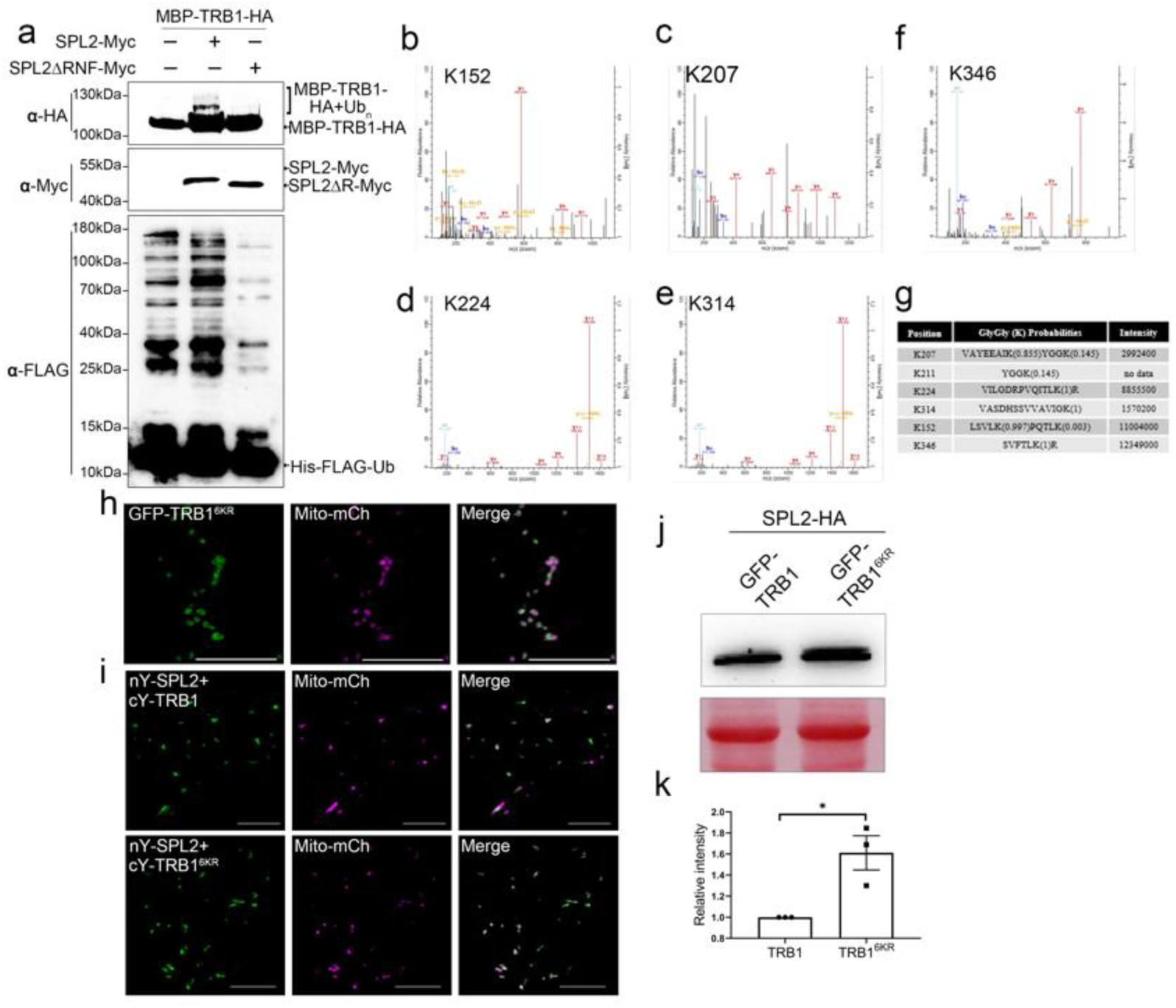
The identification of the TRB1 ubiquitination site. **a.** SPL2 mediates the ubiquitination of TRB1 *in vitro*. Bacterial lysates from *E. coli* strains expressing AtUBA1-S, MBP-TRB1-HA, AtUBC8-S, His-FLAG-UBQ10, and either SPL2-Myc or SPL2ΔR-Myc were analysed by Western blotting with an anti-HA antibody to detect TRB1 ubiquitination. As controls, SPL2 protein expression was assessed using an anti-Myc antibody. An anti-FLAG antibody was used to detect FLAG-ubiquitin (Ub) conjugates. **b-g.** Liquid chromatography-tandem mass spectrometry (LC–MS/MS) analysis was performed to identify putative TRB1 ubiquitination sites *in vivo*. The results indicate TRB1 ubiquitination at residues K152 (b), K207 (c), K224 (d), K314 (e), and K346 (f). **g.** Table of all predicted TRB1 ubiquitination sites. Among the five identified lysine residues, a new potential ubiquitination site (K211) was predicted using http://ubpred.org/. **h.** *N. benthamiana* cells co-expressing GFP-TRB1^6KR^ and Mito-mCherry indicate that the mutated TRB1 still localises to the mitochondrial outer membrane. Scale bar, 10 μm. **i.** The Split-YFP BiFC assay showed that nYFP-SPL2 and cYFP-TRB1^6KR^ produced fluorescent signals at the mitochondrial outer membrane when co-expressed in *N. benthamiana*, indicating that TRB1^6KR^ and SPL2 still interact. Scale bar, 10 μm. **j-k.** Compared with TRB1, TRB1^6KR^ levels are higher in the presence of SPL2 in *N. benthamiana*. Three biological replicates were used for quantification. Error bars represent SEM, and asterisks denote statistically significant differences in means at *P* < 0.05 (two-tailed Student’s *t*-test).

**Extended Data Fig. 5.**
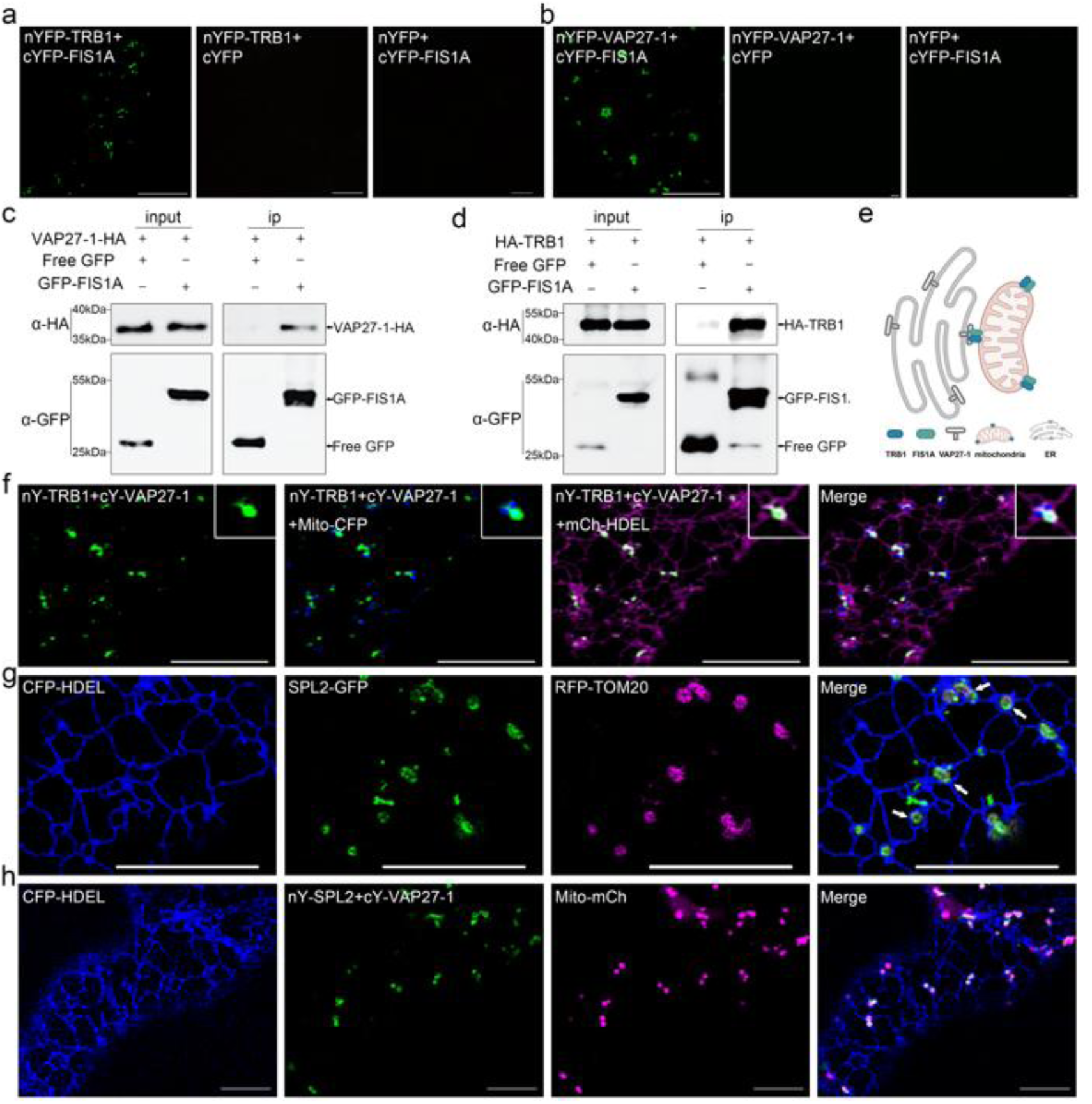
The TRB1-FIS1A-VAP27-1 complex localises to mitochondria-ER contact sites (EMCSs), with SPL2 showing partial EMCS association. **a-b.** BiFC assays of cYFP-FIS1A and nYFP-TRB1(a) or cYFP-FIS1A and nYFP-VAP27-1 (b) demonstrate that FIS1A interacts with both TRB1 and VAP27-1. **c-d.** GFP-Trap assay demonstrates GFP-FIS1A interaction with both HA-TRB1 and VAP27-1-HA in *N. benthamiana* leaf epidermal cells. VAP27-1-HA (c) and HA-TRB1 (d) are pulled down only in the presence of GFP-FIA1A (right) but not with free GFP (left). Proteins were extracted 48 h post-infiltration. **e.** Putative model of TRB1-FIS1A-VAP27-1 complex at the mitochondria-ER contact sites. **f.** The VAP27-TRB1 BiFC signals colocalised with Mito-CFP and mCherry-HDEL, demonstrating that the interaction occurs at mitochondria-ER contact sites. **g.** The co-expression of CFP-HDEL, SPL2-GFP, and RFP-TOM20-3 in *N. benthamiana*. White arrow indicates SPL2 closely associated with the ER. **h.** BiFC assay indicates an interaction between VAP27-1 and SPL2 in *N. benthamiana*. The YFP signal partially colocalised with CFP-HDEL and Mito-mCherry, demonstrating that the SPL2-VAP27-1 interacts at the ER-mitochondrial interface. Scale bar = 10 µm in confocal images.

**Extended Data Fig. 6.**
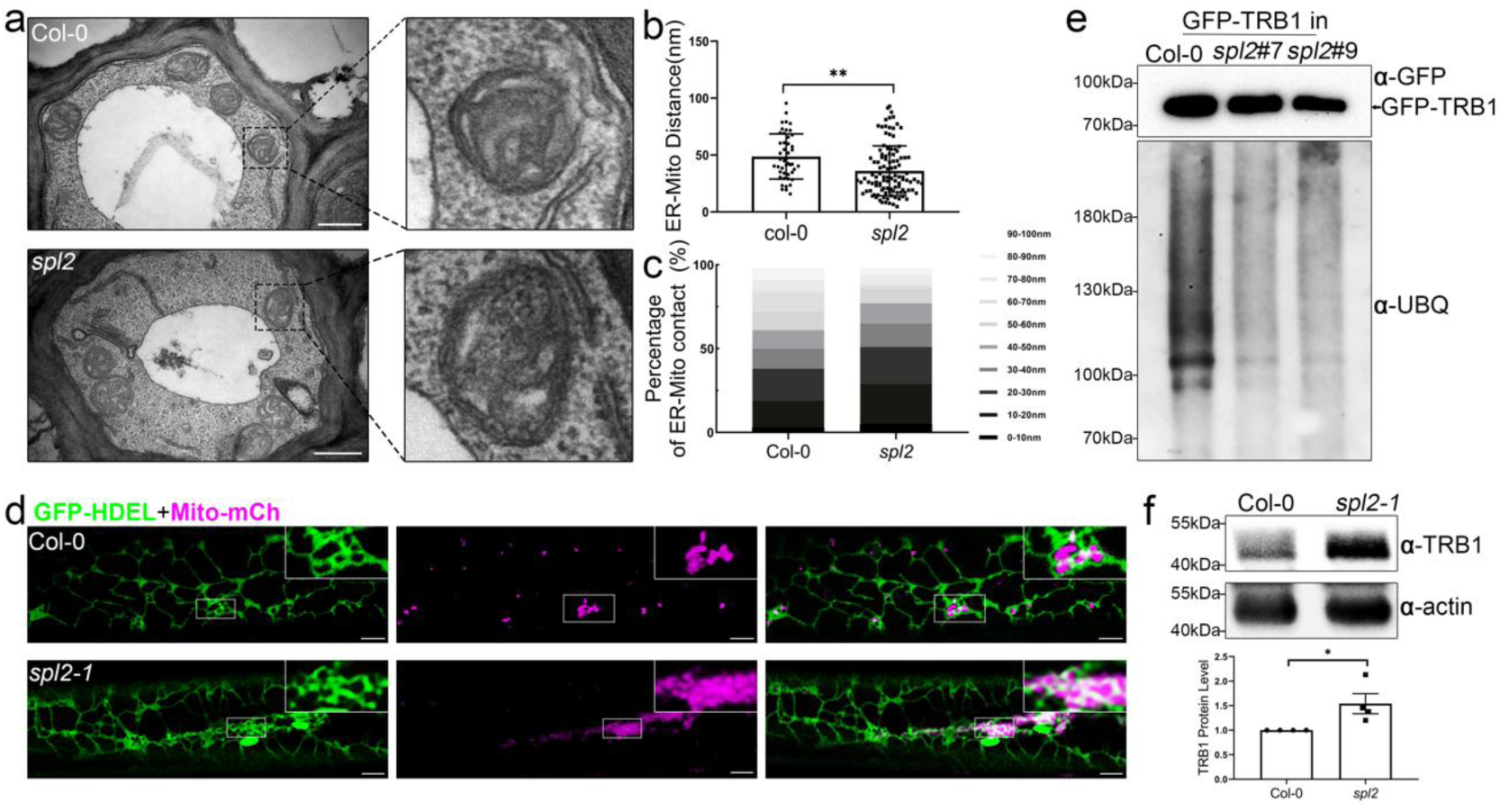
The s*pl2* mutants exhibit reduced ER-mitochondria distance and TRB1 protein accumulation. **a-c.** TEM study of wild-type Arabidopsis and *spl2-1* mutant, the distance between ER and mitochondria is significantly reduced compared to Col-0. At least 80 mitochondria from 40 cells of over three independent experiments were examined. Scale bar = 500 nm. **d.** Images highlight ER membranes closely opposed to mitochondrial aggregates in 5-day-old etiolated *spl2-1* hypocotyl cells, as well as abnormal ER aggregates. Scale bar = 10 µm**. e.** TRB1 ubiquitination is reduced in *spl2-1* mutant plants. Stable transgenic Arabidopsis lines (35S::GFP-TRB1) were generated and crossed into the *spl2-1* mutant background. GFP-TRB1 was subsequently purified from plants using GFP-Trap beads for comparative analyses. **f.** The level of endogenous TRB1 (using a TRB1 antibody) is higher in the *spl2-1* mutant than in the wild-type plants. Protein levels were quantified relative to actin (loading control), with the Col-0 level normalized to 1 in each biological replicate. Three biological repeats were taken for the quantification. Error bars represent SEM; asterisks indicate significant differences (*P* < 0.05, two-tailed Student’s *t*-test).

**Extended Data Fig. 7.**
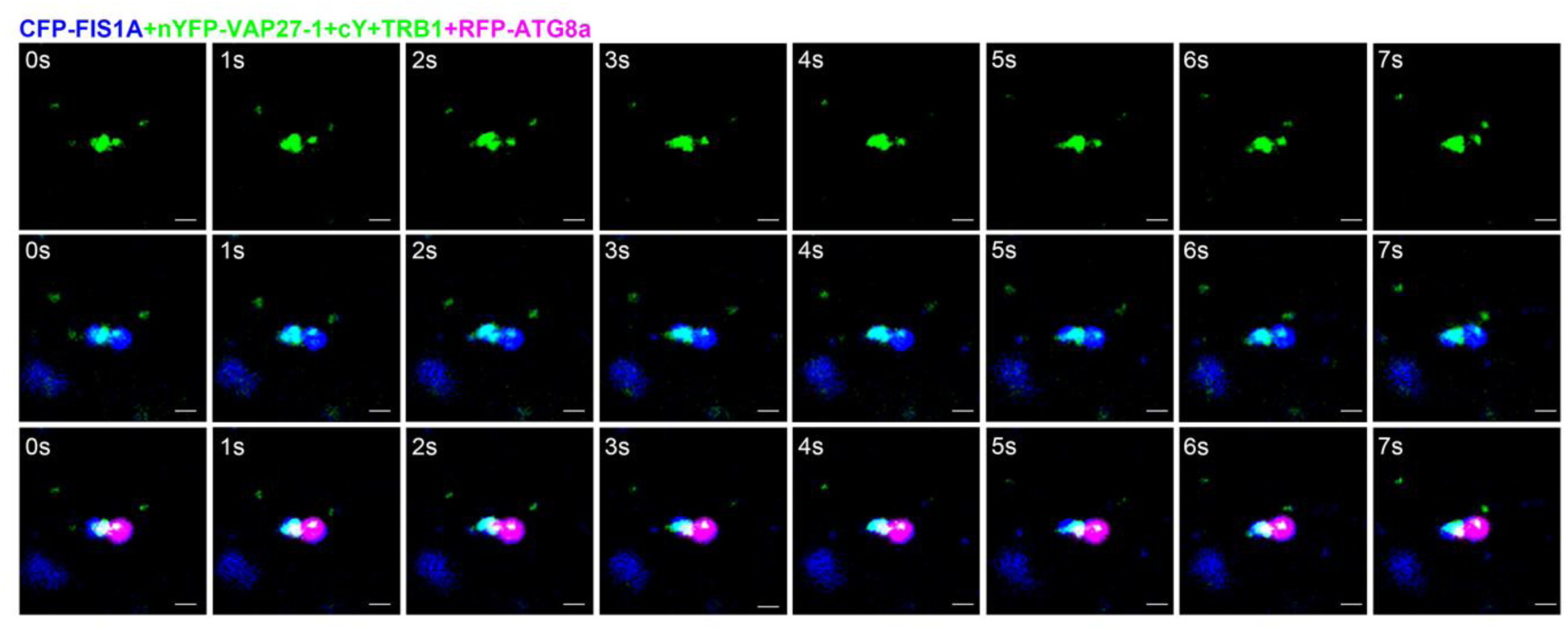
Time-lapse imaging of FIS1A-TRB1-VAP27-1-labelled ER-Mitochondrial contact sites and their partial engulfment by ATG8a-labelled autophagosomes. Time-lapse imaging of CFP-FIS1A (blue), nYFP-VAP27-1 and cYFP-TRB1 (green), and RFP-ATG8a (magenta) in *N. benthamiana*. FIS1A-labelled mitochondria (blue) co-localised with TRB1-VAP27-1 labelled ER-mitochondrial contact sites (green), and both structures are engulfed by RFP-ATG8a-labelled autophagosomes (magenta). Scale bar, 1 μm.

**Extended Data Fig. 8.**
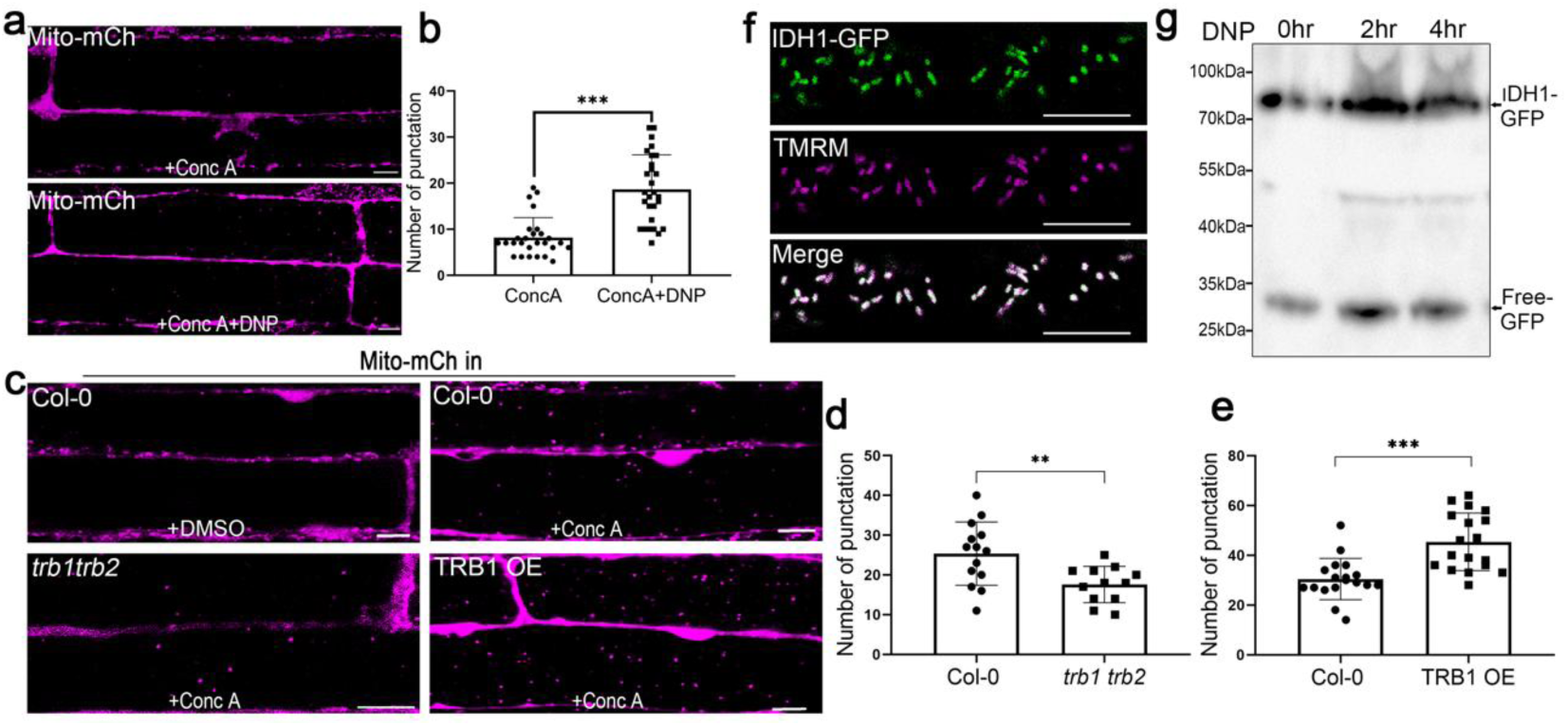
Mitophagy is stimulated by either DNP treatment or TRB1 overexpression. **a-b.** DNP treatment leads to the accumulation of mitochondria in the vacuoles in 5-day-old hypocotyl cells, indicative of active mitophagy. Scale bar, 10 μm. **c**. Representative images of 5-day-old Arabidopsis seedlings expressing Mito-mCherry in Col-0, *trb1 trb2* double mutant, and TRB1-overexpressing (OE) lines after ConcA treatment. Scale bar, 10 μm. **d-e.** Quantification of vacuole-accumulated mitochondria in the *trb1 trb2* double mutant (d) and TRB1-overexpressing lines (e) after Conc A treatment. TRB1-OE lines exhibit enhanced mitochondrial accumulation in vacuoles compared to Col-0, whereas mitochondrial degradation is suppressed in *trb1 trb2* double mutants. DMSO-treated cells were used as the control. **f-g.** Mitochondrial localised IDH1-GFP serves as a reporter for DNP-induced mitophagy. **f.** Representative images showing colocalization of IDH1-GFP (green) and mitochondria (TMRM, magenta) in root cells. Scale bar, 10 μm. **g.** The generated IDH1-GFP transgenic plants are responsive to DNP-induced mitophagy. Error bars represent SD, and asterisks indicate statistically significant differences (*P* < 0.05) as determined by a two-tailed Student’s *t*-test for b, d, e.

**Extended Data Fig. 9.**
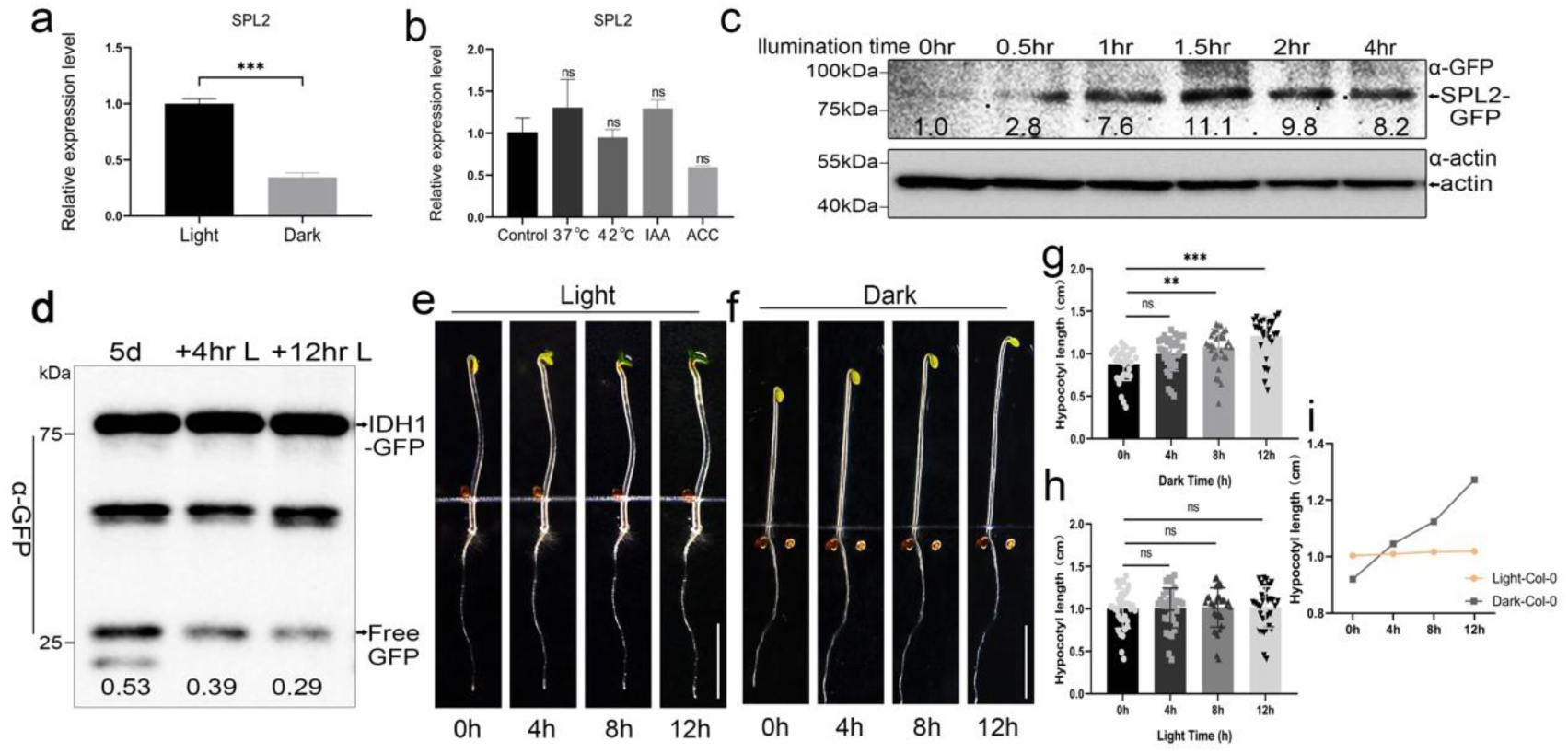
SPL2 expression is downregulated in the dark, a change associated with enhanced rapid hypocotyl elongation. **a-b.** RT-qPCR analysis reveals significantly lower *SPL2* transcript levels in dark-grown seedlings compared to light-grown controls (*** *P* < 0.001, Student’s *t*-test). In comparison, the *SPL2* expression exhibits little variation at other conditions (n.s., *P* > 0.05). **c.** Immunoblot analysis of SPL2 protein level of 5-day-old seedlings (pSPL2::SPL2-GFP/*spl2*) during de-etiolation. Actin was used as a loading control. **d.** GFP cleavage assay reveals a reduction of mitophagy flux during de-etiolation (decreased free GFP / IDH1-GFP protein ratio). Immunoblot analysis of the IDH1-GFP transgenic line: seedlings were grown on 1/2 MS medium in darkness and then transferred to a light condition at the indicated times (0, 4, 12 h). **e-f.** Hypocotyl growth ceases after light exposure. Three-day-old etiolated seedlings (Col-0) were transferred to light and imaged at the indicated time point, while dark-grown seedlings served as the control. Scale bar, 0.5 cm. **g-i.** Quantification of hypocotyl length of 3-day-old etiolated seedlings under dark (f, g) or light (e, h) conditions for 12 hours (n =30).

**Extended Data Fig. 10.**
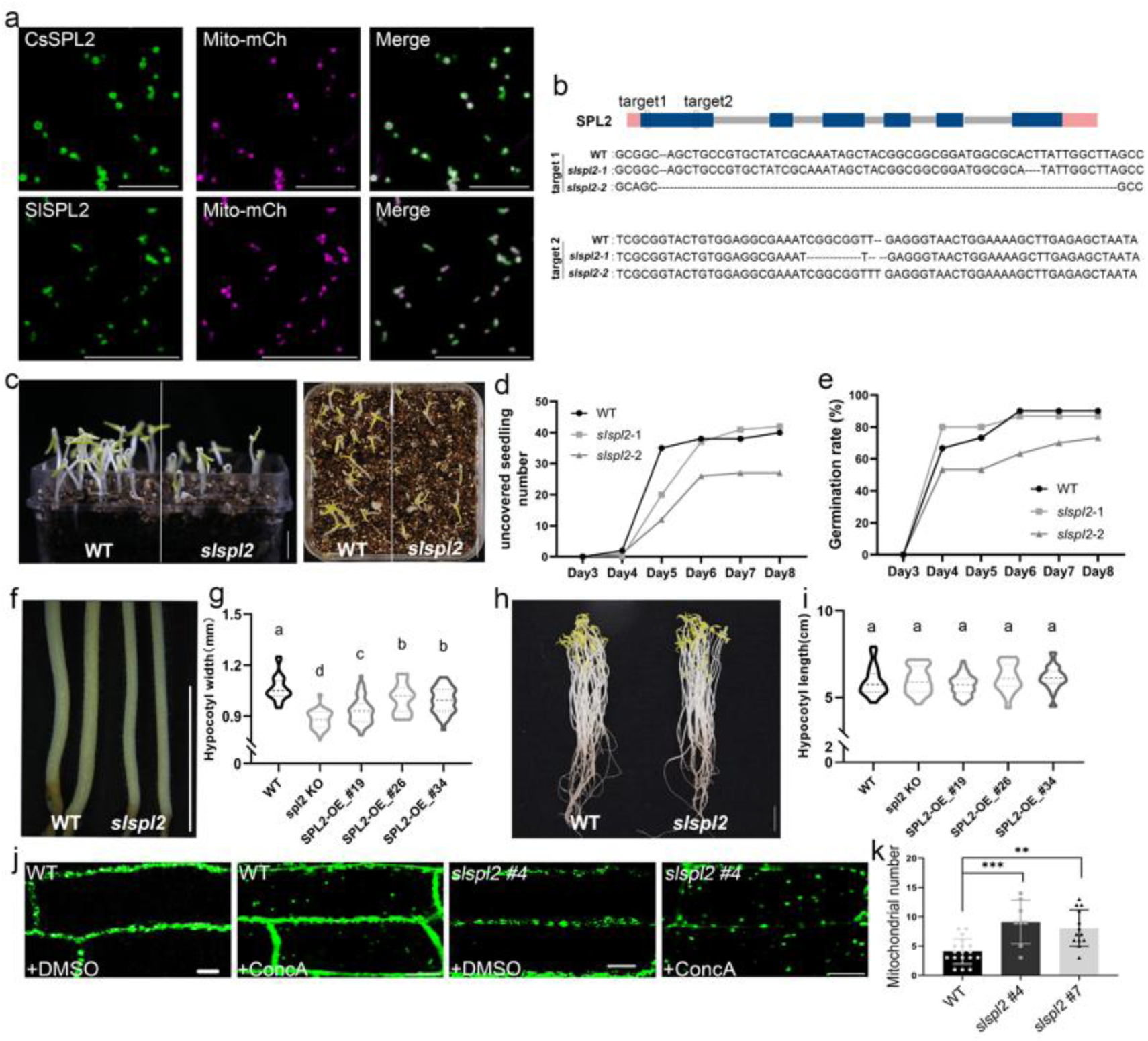
SPL2 is evolutionarily conserved in regulating mitophagy and seedling soil emergence. **a.** Co-expression of SlSPL2-GFP (tomato) or CsSPL2-GFP (citrus) with Mito-mCherry in *N. benthamiana* demonstrates mitochondrial localisation of both SPL2 orthologs. Scale bar, 10 μm. **b.** Schematic diagram of SlSPL2 genes. The sequences of the selected targets are shown below the diagram. Two homozygous double mutants, *slspl2-1* and *slspl2-2*, were obtained with deletions at the target sites. **c-d.** The *slspl2* mutant exhibited delayed seedling emergence compared to wild-type (WT) seeds. To quantify emergence time, fifty seeds each of WT and *slspl2* mutants were covered with a 1 cm layer of soil. Trays were sealed with parafilm and incubated in complete darkness. The number of seedlings penetrating the soil surface was recorded daily. Scale bar, 1 cm. **e.** Germination rates of wild-type and *slspl2* seeds. **f-i.** Hypocotyls of *slspl2* mutant seedlings grown in complete darkness for 7 days were significantly thinner than those of wild-type (WT) controls, while hypocotyl length showed no significant difference between genotypes under these conditions. Different letters represent statistically different means. Scale bar, 1 cm. **j.** The mitophagy activity in the *slspl2* plant is significantly affected. Representative confocal micrographs of roots from eight-day-old tomato seedlings. Wild-type plants and *slspl2* plants expressing the mitochondrial marker GFP-TOM20-3 were cultivated on 1/2 MS growth medium in darkness for 8 days. Following treatment with 1 µM Conc A, a greater accumulation of mitochondria within the vacuole was observed in *slspl2* seedlings compared to WT. Little vacuolar GFP signal was detected in DMSO-treated control cells of either genotype (Scale bar = 10 µm). **k.** Quantification of (j), the number of mitochondria accumulated within the vacuole per field (80µm x 40 µm) increased in *slspl2* cells after Conc A treatment.

